# Fibro-Adipogenic Progenitors require autocrine IGF-I in homeostatic and regenerating skeletal muscle

**DOI:** 10.1101/2025.04.11.648330

**Authors:** Yangyi E. Luo, Zoe Abe-Teh, Tarek Y. Alsaghir, Li-Ying Kuo, Fahong Yu, Bradley E. Stoker, Ambili Bai Appu, Yumei Zhou, Feng Yue, Daniel Kopinke, Elisabeth R. Barton

## Abstract

Fibro-Adipogenic Progenitors (FAPs) are mesenchymal stem cells that are vital for muscle homeostasis and regeneration but produce fibrosis and intramuscular fat under pathological conditions. Insulin-like Growth Factor-I (IGF-I) is a key regulator of muscle repair, satellite cell activity, macrophage polarization, and extracellular matrix (ECM) remodeling. We generated inducible FAP-specific *Igf1* deficient (FID) mice to determine the necessity of FAP IGF-I. After BaCl_2_ injury, FID mice exhibited impaired muscle regeneration, with fewer Pax7+ cells, increased macrophage accumulation, smaller fibers, reduced ECM, and depressed FAP proliferation. Following glycerol injury, FID muscles exhibited reduced adipocyte accumulation. Primary FAPs isolated from injured FID muscles had blunted growth, upregulation of immune-regulatory genes and downregulation of ECM and cell proliferation genes, with delayed responses to fibrogenic and to adipogenic media. FAP property alterations were already present in homeostatic muscle, indicated by scRNASeq, with decreased indices of protein translation and ECM production as well as increased markers of senescence, confirmed *in vivo* and *in vitro*. Overall, FAP IGF-I is a critical autocrine factor, with further paracrine consequences for muscle regenerative capacity.

## Introduction

Skeletal muscle comprises approximately 40% of human body mass and has extraordinary regenerative capacity(Bodine *et al*, 2023). Skeletal muscle regeneration is a period of dynamic remodeling that requires coordinated interaction among multiple cell types, including satellite cells (SCs), immune cells, and Fibro-Adipogenic Progenitor cells (FAPs), to restore functional muscle fibers supported by a healthy extracellular matrix (Caballero-Sanchez *et al*, 2024; Kuang *et al*, 2007; Relaix *et al*, 2005; Tidball, 2005). FAPs are mesenchymal progenitor cells that reside outside of the basal lamina surrounding skeletal muscle fibers that undergo rapid proliferation upon acute injury, and they are required for skeletal muscle regeneration and homeostasis (Collins & Kardon, 2021; Flores-Opazo *et al*, 2024; Joe *et al*, 2010; Loomis & Smith, 2023; Wosczyna *et al*, 2019). Without FAPs, skeletal muscle is subject to progressive muscle loss and delayed regeneration(Wosczyna *et al*., 2019). Although FAPs are beneficial for muscle regeneration and physiological function, they are also the primary progenitor cells responsible for excessive fibrosis and intramuscular fat deposition under pathological conditions(Kopinke *et al*, 2017; Lemos *et al*, 2015; Uezumi *et al*, 2010). Studies over the past 15 years have established that FAPs are essential in skeletal muscle biology, yet the mechanisms of how FAPs interact and regulate cell function in muscle regeneration has not been fully elucidated.

In addition to multiple cell types, many secreted proteins coordinate the process of muscle regeneration (Forcina *et al*, 2020). The interplay of myokines secreted from and acting on multiple cell types within muscle is critical for skeletal muscle remodeling. One such protein is Insulin-like Growth Factor-I (IGF-I), which facilitates skeletal muscle hypertrophy, muscle regeneration, and glucose regulation (Barton-Davis *et al*, 1998; Luo *et al*, 2024; Musaro *et al*, 2001; Vassilakos *et al*, 2019). A major source of IGF-I is the muscle fiber, which produces up to 75% of the local IGF-I pool (Barton *et al*, 2012). Prior studies from our group demonstrated that IGF-I from muscle fibers is important for post-natal growth and functional recovery from disuse atrophy (Spradlin *et al*, 2021; Vassilakos *et al*., 2019). However, other IGF-I sources also have impact on muscle regenerative capacity, including myeloid cells where loss of *Igf1* expression impairs macrophage polarization and ultimately slows regeneration (Luo *et al*., 2024; Tonkin *et al*, 2015). The contribution of IGF-I from FAPs has not been evaluated, even though single cell RNAseq databases implicate FAPs as having one of the highest *Igf1* expression levels throughout muscle regeneration (Kok & Barton, 2021). Therefore, it is important to determine if IGF-I from FAPs is a significant contributor to FAP behavior, and by extension, FAP actions on muscle physiology, pathology, and intercellular interactions under multiple conditions(Kim *et al*, 2022; Uezumi *et al*., 2010; Wosczyna *et al*., 2019). In this study, we hypothesize that IGF-I secreted from FAPs is critical for their function. We investigated the significance of IGF-I secreted from FAPs on skeletal muscle properties, including regenerative capacity, as well as FAP proliferative and differentiation capacity, using a mouse model that afforded FAP specific deletion of *Igf1*. This study provides multiple lines of evidence that IGF-I produced by FAPs is an important autocrine factor for FAP actions during muscle regeneration. The consequences of its loss alter FAPs to an extent that they are essentially removed from performing their normal interactions with other cell populations in the muscle, and ultimately stall regenerative capacity.

## Results

### Deletion of IGF-I in PGDFRα+ mesenchymal progenitor cells in skeletal muscle moderately reduces IGF-I pool

FAPs are muscle resident mesenchymal progenitor cells marked by their distinct surface protein Platelet-Derived Growth Factor Receptor Alpha (PDGFRɑ) (Contreras *et al*, 2019; Kopinke *et al*., 2017; Mueller *et al*, 2016; Norris *et al*, 2023; Uezumi *et al*., 2010; Wosczyna *et al*., 2019). Although it has been documented that FAPs secrete IGF-I during muscle homeostasis and regeneration, the significance of IGF-I from FAPs is not clear. Thus, we generated a conditional FAP-specific knockout mouse line named <u>F</u>AP-IGF-I-<u>D</u>eficient (FID) by breeding inducible *Pdgfrα*^Cre-ERT2^ mice (Chung *et al*, 2018) with *Igf1*^fl/fl^ mice (Spradlin *et al*., 2021) and the Cre-dependent lineage reporter mouse *Rosa26*^EYFP^ mice (Srinivas *et al*, 2001) (Figure 1A). Since the insertion of *Pdgfrɑ*^CRE-ERT2^ allele does not result in observable phenotypic changes (Figure S1E-H), animals without the *Pdgfrɑ*^CRE-ERT2^ allele served as control groups (*Igf1^fl/fl^*) (Figure 1A).

**Figure 1.**
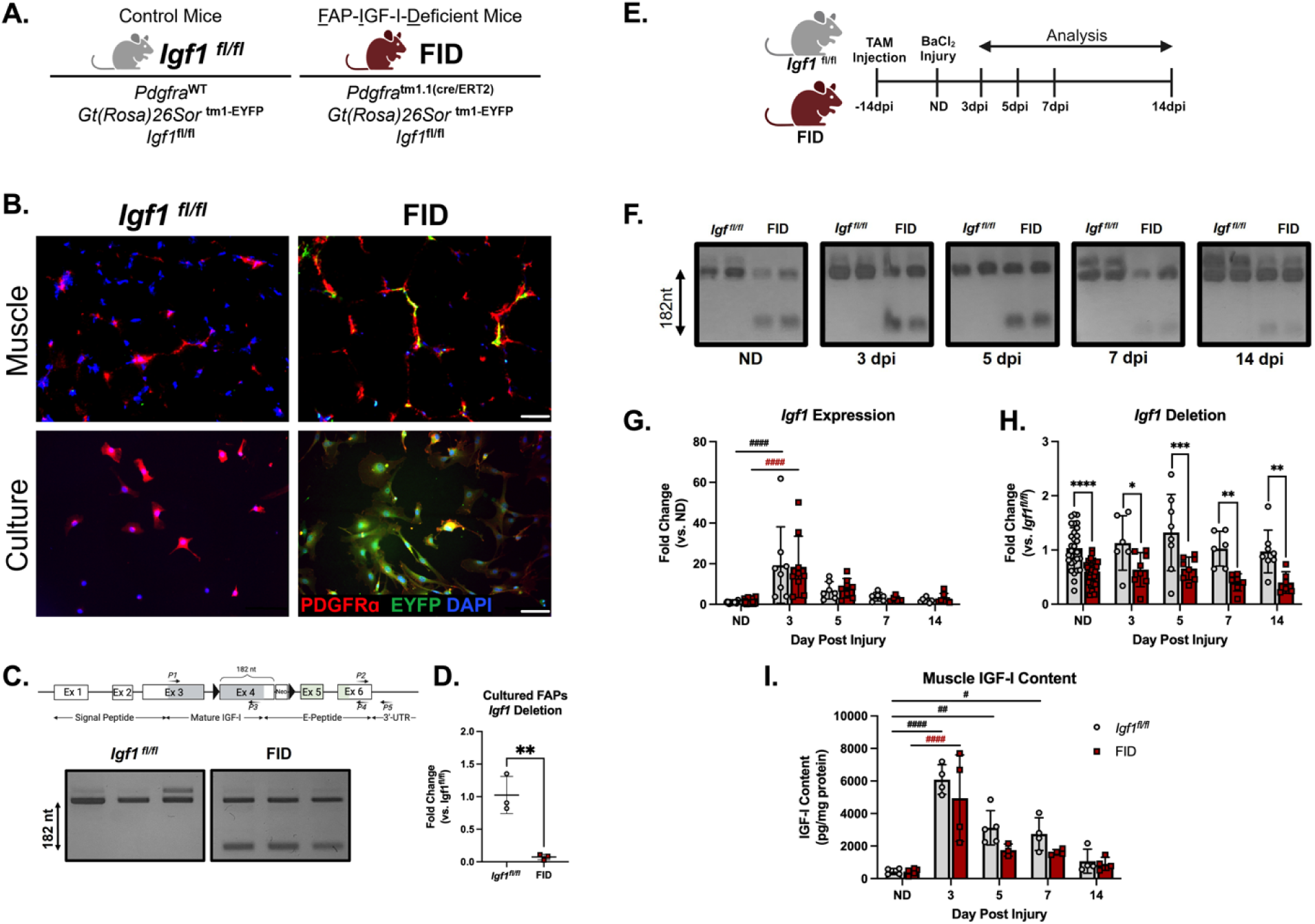
Validation of FID mouse model and the specificity and efficiency of *Igf1* deletion. A: Genetic scheme of control (*Pdgfrα*^WT^x *Igf1^fl/fl^* x *Rosa26*^EYFP^) and FID (*Pdgfrα*^CRE-ERT2^ x *Igf1^fl/fl^* x *Rosa26*^EYFP^) mouse models. B: Immunofluorescent staining of PDGFRa^+^(red), GFP (green), and nuclei by DAPI (blue) on primary FAPs and muscle cryosections of non-damaged TA muscle. Scale Bar: 200µm. C: Igf1 exons shown with the location of primers (P) for expression analysis. Endpoint RT-PCR examination of *Igf1* exon 4 deletion (primers P1 and P4) in primary FAPs isolated from 3dpi. The upper band shows amplification of *Igf1* cDNA from exon3 to exon 6 and the lower bands is the result of exon 4 excision by Cre-recombinase. D: qRT-PCR analysis of *Igf1* deletion using primers P1 and P3 targeting exon 3 and exon 4 (region of deletion) and normalized against total *Igf1* transcripts (primers P2 and P5, Exon 6-3’-UTR) in cultured primary FAPs. p=0.0047 by unpaired t-test. E: Illustration of the acute muscle regeneration experimental design. F: Endpoint RT-PCR examination of *Igf1* exon 4 deletion (primers P1 and P4) in TA muscles throughout regeneration. The upper bands show amplification of *Igf1* cDNA from exon3 to exon 6 and the lower band is the result of exon 4 excision by Cre-recombinase. G: qRT-PCR using primers P2 and P5 targeting the exon 6 to 3’ UTR region of the *Igf1* shows there is no change in endogenous *Igf1* expression between control and FID muscles, and that there is a transient increase in *Igf1* expression during regeneration. Two-way ANOVA: Interaction=ns, timepoints p<0.0001, strains=ns. n=19 for control and n=25 for FID at ND, n=6∼8 for both strains from 3dpi to 14dpi. H: qRT-PCR analysis of *Igf1* deletion using primers P1 and P3 targeting exon 3 and exon 4 (region of deletion) and normalized against total *Igf1* transcripts (primers P2 and P5, Exon 6-3’-UTR). There is significant *Igf1* deletion at all times of interest during homeostasis and regeneration. Two-way ANOVA followed by Šídák post-hoc test: ND p= 0.000002, 3dpi p=0.044, 5dpi p=0.023, 7dpi p=0.0008, 14dpi p=0.007. n=19 for control and n=25 for FID at ND, n=6∼8 for both strains from 3dpi to 14dpi. I: ELISA analysis of muscle tissue IGF-I protein levels during acute muscle regeneration. IGF-I produced from FAPs contributes about 20% to the total IGF-I content in TA muscle throughout regeneration. Two-way ANOVA: interaction=ns, timepoints p<0.0001, strains p=0.0238. n=3∼6 for both strains from ND to 14dpi. Two-way ANOVA (strains and injury time) followed by Šídák post-hoc test used for statistical analysis. # represents comparison between timepoints and * represents comparison between strains. * or # p<0.05, ** or ## p<0.01, *** or ### p<0.001, **** or #### p<0.0001.

Confirmation of efficient and cell specific Cre expression was confirmed *in vivo* and *in vitro*. The colocalization of PDGFRɑ antibody staining and the EYFP reporter in tissue sections was evident after tamoxifen (TAM) treatment with no reporter signal observed without co-labeling by PDGFRɑ (Figure 1B). Similar labeling patterns were observed in primary FAPs isolated from TAM treated mice 3 days after acute injury (Figure 1B). To determine *Igf1* deletion efficiency, endpoint RT-PCR was utilized with primers flanking the floxed exon 4 as previously performed (Figure 1C, primers P1 and P4) (Vassilakos *et al*., 2019), in FAPs isolated from FID and *Igf1^fl/fl^*muscles 3 days post injury (dpi). Only FID FAPs displayed amplification of a smaller band indicative excision of exon 4, with a faint doublet observed in FAPs from *Igf1^fl/fl^* muscles originating from alternative splicing inclusion of exon 5 (Figure 1C). Next, qRT-PCR amplification by primers targeting exons 3 and 4 (primers P1 and P3) was normalized to those amplifying exon 6 and 3’UTR (primers P2 and P5) to quantify the proportion of transcripts retaining exon 4. Deletion of exon 4 occurred in at least 80% of the transcripts in cultured FAPs (Figure 1D).

There are multiple sources of IGF-I in skeletal muscle (Kok & Barton, 2021), therefore we examined the impact of FAP *Igf1* ablation on gene expression and protein production of IGF-I in homeostatic muscle and throughout regeneration. Muscle injury was induced by 1.2% BaCl_2_ delivered via intramuscular injections into the Tibialis Anterior (TA) muscle, and tissues were analyzed from a non-damaged (ND) state through 14 days post injury (14 dpi) (Figure 1E). Endpoint RT-PCR demonstrated that exon 4 excision was evident only in FID muscles both in ND and following injury (Figure 1F). Because *Igf1* expression changes dramatically throughout regeneration (Luo *et al*., 2024), we determined if the general regulation of *Igf1* transcription was altered by the ablation of FAP *Igf1*. We measured all *Igf1* transcripts by qRT-PCR, using primers that targeted exon 6 and the 3’-UTR sequences shared by all *Igf1* isoforms and unaffected by the deletion of exon 4 (primers P2 and P5, Figure 1C) (Barton, 2006). Total *Igf1* expression was significantly elevated in muscles 3 days post injury (3dpi) compared to ND muscles, but the transient increase of *Igf1* expression was not affected by strain (Figure 1G). Next, extent of deletion was measured in regenerating muscles following the same strategy used in cultured FAPs (Figure 1D). There was a 20-45% reduction in functional *Igf1* transcripts in regenerating FID muscles that persisted throughout regeneration (Figure 1H). Lastly, IGF-I protein was analyzed in regenerating muscles from *Igf1^fl/fl^*and FID mice. The transient >10-fold increase in IGF-I content was apparent in both strains, with the difference between *Igf1^fl/fl^* and FID muscles leading to significance for both timepoint and strain during regeneration (Figure 1I). However, the contribution of 20-30% IGF-I from FAPs at any given timepoint did not achieve significance with post-hoc testing. In sum, the cell specific *Igf1* deletion strategy was successful in removing the production of IGF-I from FAPs in homeostatic and regenerating muscle. The contribution of FAP IGF-I to the total local pool is approximately 25%, in comparison to other local sources, such as muscle fibers and myeloid cells (Tonkin *et al*., 2015; Vassilakos *et al*., 2019).

In previous studies, we found that deletion of *Igf1* from muscle fibers in adult mice altered anabolic and metabolic properties (Spradlin *et al*., 2021; Vassilakos *et al*., 2019). Further, deletion of FAPs from skeletal muscle also caused rapid loss of muscle mass in adult mice (Uezumi *et al*, 2021; Wosczyna *et al*., 2019). To determine if the FAP source of IGF-I affected the muscle phenotype, muscles were examined up to 4 weeks after TAM induction of deletion. There were no changes in muscle mass, fiber type or fiber size in muscles from FID mice compared to *Igf1^fl/fl^* mice (Figure S1C-D, 2D-F). Thus, the 25% reduction of total IGF-I in FID muscles did not appear to alter muscle properties, and it is not essential for their maintenance.

### IGF-I from FAPs is required for early and late stages of muscle regeneration

FAPs are essential for muscle regeneration (Wosczyna *et al*., 2019). Therefore, we investigated if IGF-I from FAPs was critical for muscle regeneration using muscles from mice after intramuscular TA injections of BaCl_2_ (Figure 2A). Even though no differences in muscle masses or fiber sizes in ND muscles were apparent post-TAM injection (Figure 2D-F, S1C-D), regenerating muscles displayed FID-dependent differences in recovery of mass. Specifically, we observed a significant reduction in relative muscle mass (damaged LTA/ND RTA) and median fiber size at 7dpi in both male and female FID mice compared to age and sex-matched controls; all groups returned to ND mass and fiber size by 14 dpi (Figure 2B-H). This suggests that FAPs produce a significant IGF-I source as FAPs undergo proliferation (Joe *et al*., 2010; Kopinke *et al*., 2017) early after injury, and that it is important for the efficiency of early stages of regeneration. We extended the analysis of regenerating muscles to 28 dpi, where we found a second point at which muscles from FID mice were significantly smaller than those from *Igf1^fl/fl^* mice. FAP muscle fiber size plateaued after 14 days and failed to grow, in contrast to those from WT mice. Hence, the effect of IGF-I loss from FAPs in biphasic, with both early and late-stage deficits in regeneration, in male and female FID mice.

**Figure 2.**
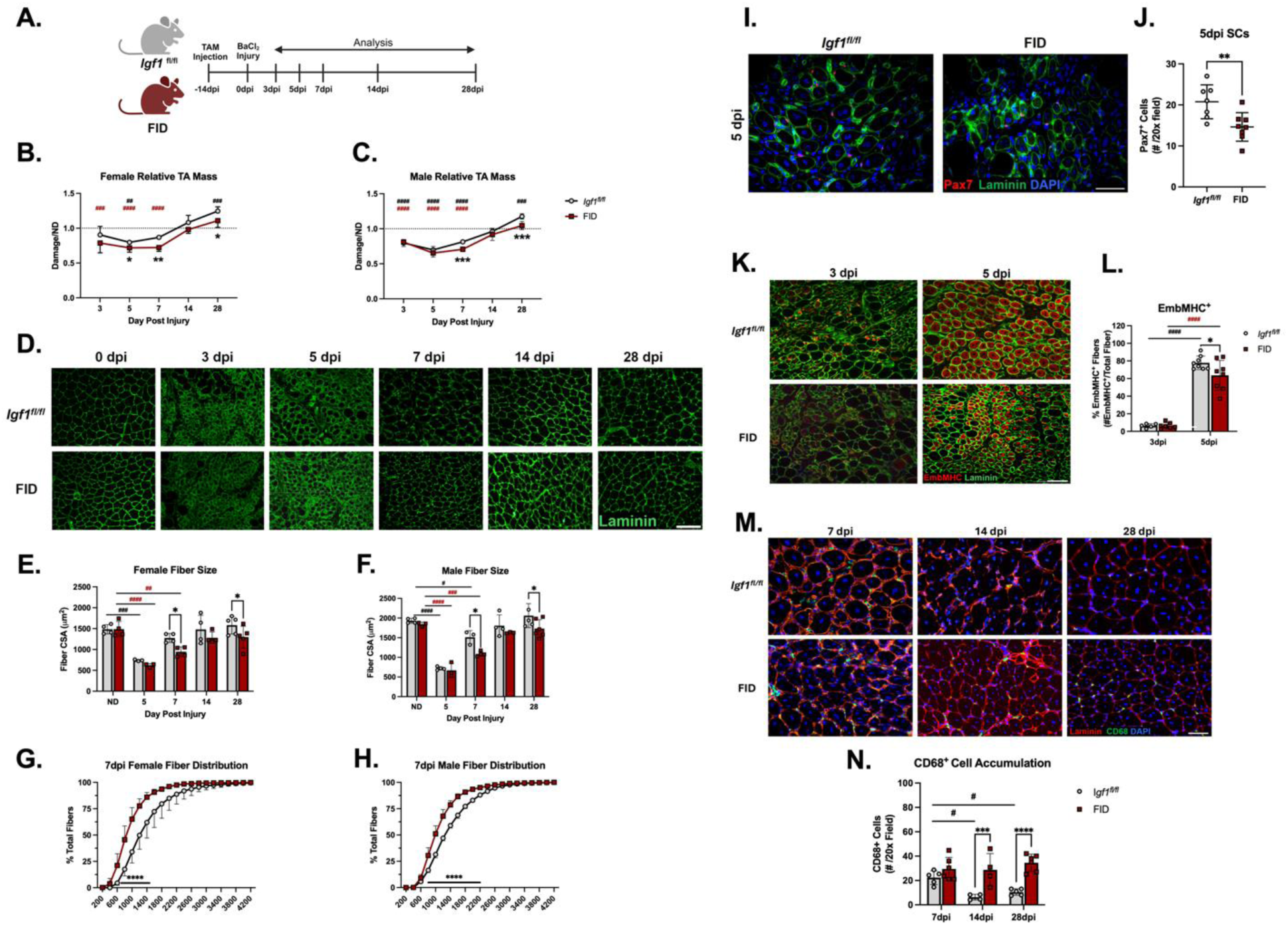
IGF-I produced by FAPs is required for early muscle regeneration. A: Illustration of the acute muscle regeneration study design. B-C: Relative muscle weight in both females and males shows decrease in muscle mass during regeneration that is exacerbated in FID muscles at 7 and 28 dpi. FID muscles from both sexes showed a significant reduction in relative muscle mass at 3 – 7 dpi. *Igf1^fl/fl^* muscles from both sexes showed a significant increase in relative muscle mass at 28 dpi. The muscle relative mass is obtained by normalizing the damaged LTA to the non-damaged RTA. Female two-way ANOVA: interactions p=ns, timepoints p<0.0001, strains p=0.0011. Male two-way ANOVA: Interactions p=ns, timepoints p<0.0001, strains p=0.0244. N=3∼4 for both strains and sexes from 3dpi to 28dpi. D: Immunofluorescent staining of laminin (green) on muscle cryosections during muscle regeneration. Scale bar: 200µm. E-F: Muscle median cross-sectional fiber areas show smaller fibers in FID muscles at 7 and 28 dpi in both sexes. Muscle at 5dpi exhibits a significant reduction in muscle fiber size in both strains and sexes. Female two-way ANOVA: interactions p=ns, timepoints p<0.0001, strains p=0.0829. Male two-way ANOVA: Interactions p=ns, timepoints p<0.0001, strains p=0.0022. N=3∼4 for both strains and both sex from ND to 28 dpi. G-H: Cumulative muscle fiber size distribution at day 7 post-injury show that FID muscle fibers are smaller in both male and female mice. Female two-way ANOVA: interactions p<0.0001, timepoints p<0.0001, strains p<0.0001. Fiber size 400µm - 1600µm p<0.0001. Male two-way ANOVA: interactions p<0.0001, timepoints p<0.0001, strains p<0.0001. Fiber size 800µm - 2000µm p<0.0001. I: Immunofluorescent staining of Pax7 (red), laminin (green), and nuclei (DAPI, blue) in 5dpi muscle sections, where overlap between Pax7 and DAPI (pink) indicates a satellite cell (SC). Scale bar: 100µm. J: Quantification of SCs at 5dpi demonstrates there is a reduction of SCs in regenerating FID muscles. N=7 for control and n=8 for FID. Unpaired t-test p=0.0082. K: immunofluorescent staining of EmbMHC and laminin on muscle sections from 3 and 5 dpi. N=3∼4 for group of both sexes. Scale bar: 200µm. L: 3dpi and 5dpi EmbMHC quantification. %EmbMHC= # EmbMHC / # fibers. Two-way ANOVA: interactions p=ns, timepoints p<0.0001, strains p=0.0865. 5dpi p=0.0103. N=6∼8 for both strains. Scale bar: 100µm or 200µm. An unpaired student t-test or two-way ANOVA (strains and injury time) followed by Šídák post-hoc test was used for statistical analysis. # represent comparison between timepoints and * represents comparison between strains. * or # p<0.05, ** or ## p<0.01, *** or ### p<0.001, **** or #### p<0.0001

Muscle regeneration requires the actions of multiple cell types, including SCs and macrophages, to coordinate with FAPs and create new muscle. To determine the breadth of the effects of IGF-I from FAPs on the regeneration process, SC numbers were quantified by Pax7 immunostaining in muscle sections at 5 dpi, where they are most prevalent. There was a significant reduction of Pax7^+^ SCs in FID muscles at 5dpi compared to regenerating *Igf1^fl/fl^* muscles (Figure 2I-J), implicating the need for IGF-I from FAPs to help with SC activation and/or expansion following injury (Joe *et al*., 2010).

Expression of embryonic myosin heavy chain (EmbMHC/*Myh3*) is a key step in the formation of newly formed muscle fibers (Agarwal *et al*, 2020), and sections from muscles at 3 and 5 dpi were evaluated for the presence of EmbMHC positive fibers. There were EmbMHC^+^ fibers sparsely distributed throughout muscle sections at 3 dpi in both strains (Figure 2K-L). By 5 dpi, the proportion of EmbMHC^+^ fibers increased dramatically in both strains, but there was a significant reduction in regenerating FID muscles at this timepoint (Figure 2K-L). Even so, the transient upregulation of *Myod* and *Myogenin* expression in damaged muscle was unaffected by the loss of IGF-I from FAPs (Figure S2A-B). Finally, we examined sections from regenerating muscle for macrophage presence through immunostaining for CD68, which marks myeloid cells (Chistiakov *et al*, 2017). CD68^+^ cells were evident in muscles at 7 dpi and then diminished at 14 and 28 dpi in *Igf1^fl/fl^*muscles; however, their presence persisted in FID muscles at the later timepoints (Figure 2M-N). This finding suggests that the crosstalk between FAPs and macrophages to resolve damage was also disrupted with the loss of IGF-I from FAPs. Altogether, multiple measures of regeneration exhibited delays when IGF-I from FAPs was lost, and the timing of the deficits mirrored many of the deficits observed in the absence of FAPs (Joe *et al*., 2010; Wosczyna *et al*., 2019).

### FAP proliferation requires an autocrine source of IGF-I

Our data suggests that IGF-I from FAPs impacts regeneration during early and late stages, but it is unclear if it is directly modulating FAP activity. FAPs express IGF-I receptor (*Igf1r*) in regenerating, developing, and aging skeletal muscle (Kok & Barton, 2021), and so it is likely that they can respond to IGF-I regardless of its source.

Therefore, we examined the potential autocrine function of FAP IGF-I in regenerating muscle, focusing on the first 7 days after injury when FAPs are most abundant (Joe *et al*., 2010; Kopinke *et al*., 2017) (Figure 3A). First, we quantified the number of resident FAPs following injury by immunostaining muscle sections for laminin and PDGFRα (Figure 3B), and found that *Igf1^fl/fl^*muscles exhibited the well-documented transient expansion of FAPs that was maximal at 5 dpi (Figure 3C). In contrast, FAPs failed to expand in FID muscles and lacked the amplification of the FAP pool observed in *Igf1^fl/fl^* muscles (Figure 3B-C). To determine the cause of the diminished FAP numbers in regenerating FID muscles, muscle sections were stained for Ki67 and PDGFRα to mark proliferating FAPs (Figure 3D). There was a substantial 30% reduction in Ki67^+^ FAPs in regenerating FID muscles at 5 dpi (Figure 3E), indicating that FAP proliferation was impaired in the absence of FAP IGF-I *in vivo*. FAP proliferation defects could be due to altered environmental cues caused by diminished IGF-I, and thus via indirect feedback on FAP properties, or an effect directly imposed on FAPs. To determine which of these scenarios was true, primary FAPs were isolated from TA muscles at 3 dpi and plated to observe growth *in vitro*. In the process of plating FAPs, we first found a reduction in the number and proportion of FAPs initially isolated from FID muscles even though total number of cells did not differ (Figure S2C-E). Thus, cells were plated at the same density after isolation and were cultured in media without supplemental exogenous IGF-I to ensure robust comparisons between strains. Plated cells were allowed to grow for 1 week, with regular monitoring of cell numbers. FAPs from FID mice doubled in 1 week, whereas the FAPs isolated from *Igf1^fl/fl^*muscles expanded 7-fold in the same time period (Figure 3F-G). To determine if proliferation could be restored in FAPs isolated from FID muscles, exogenous recombinant IGF-I (rIGF, 10 nM) was provided, and Ki67 positive cells were measured 24 hours after plating. FAPs from *Igf1^fl/fl^* muscles had significantly more cells co-labeled with DAPI and Ki67, and the proliferation was also enhanced with exogenous rIGF-I and reversed by the addition of IGF-IR inhibitor AEW541 (Figure 3H-I). However, there was no response to increased rIGF-1 by FAPs from FID muscles, suggesting the detrimental effects on FAPs by the absence of autocrine IGF-I occurred much earlier in vivo. To assure that the IGF-I pathway was intact, immunoblotting for phosphorylation of Akt and ERK1/2 was performed under the same conditions. FAPs from FID muscles had less total ERK1/2 and Akt than those from *Igf1^fl/fl^*muscles, but cells from both strains responded to exogenous rIGF-I, and phosphorylation was blunted by AEW541 (Figure 3J), confirming an intact IGF-I signaling pathway. Taken together, the impact of *Igf1* loss from FAPs likely occurs prior to injury, leading to decrements in proliferation both *in vivo* and *in vitro* that are autocrine in nature, and show that IGF-I made by FAPs is indispensable for their proliferation.

**Figure 3.**
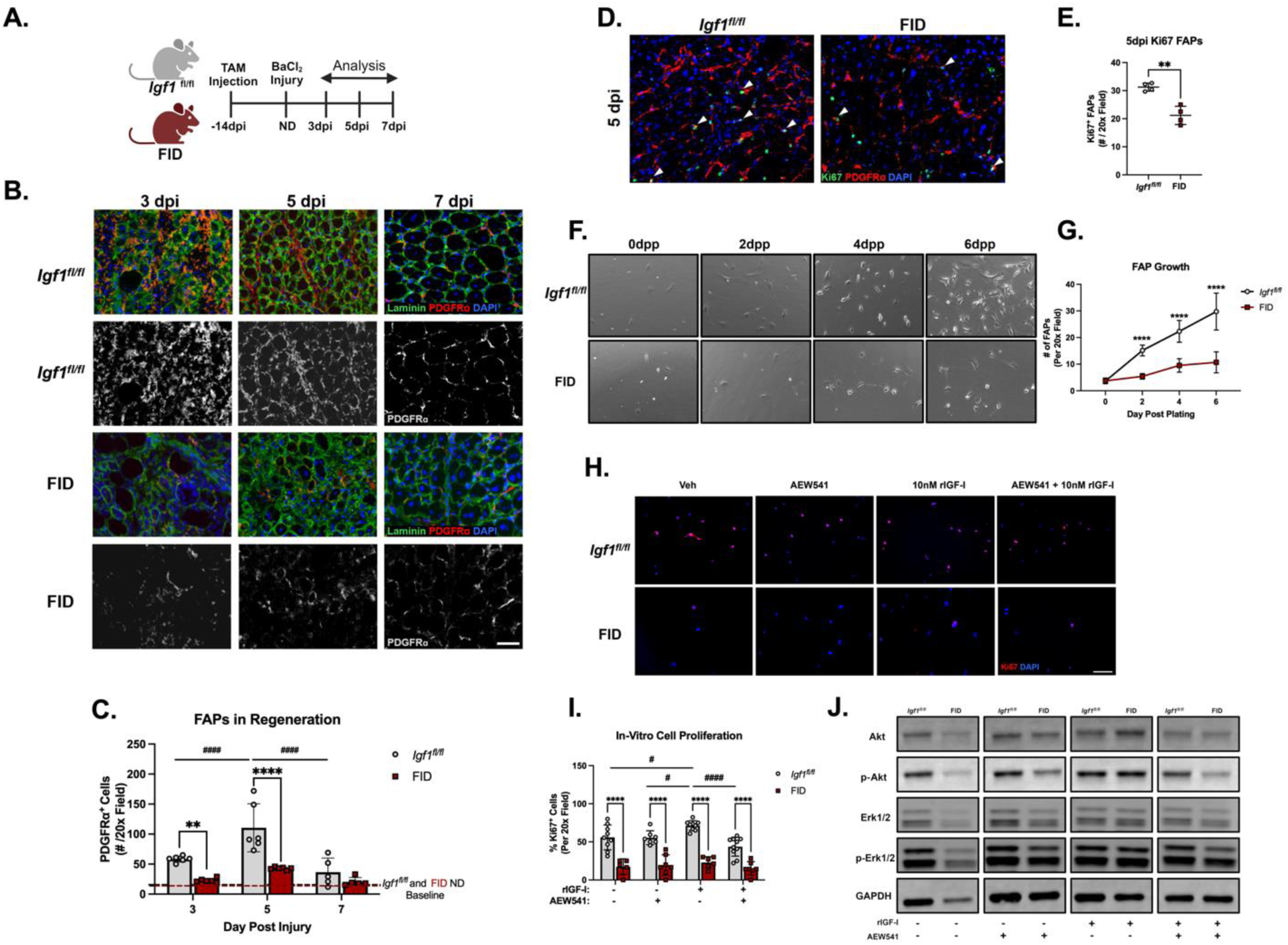
IGF-I from FAPs is required for FAPs proliferation and expansion both *in vivo* and *in vitro*. A: Graphical illustration of muscle regeneration experimental design. B: Immunofluorescent staining of PGDFRɑ (red), laminin (green), and nuclei (DAPI, blue) on sections from 3-7dpi muscles. Greyscale images on the second and fourth rows are PDGFRɑ signals converted to 8-bit images to demonstrate the difference in the number of PDGFRɑ+ cells between *Igf1^fl/fl^*and FID. Scale Bar: 100µm. C: Quantification of PDGFRɑ+ cells from 3-7dpi muscles. FAPs from *Igf1^fl/fl^*muscles showed a significant expansion from 3dpi and 5dpi, which is not being observed in FID muscles. Two-Way ANOVA: interactions p=0.014, timepoints p<0.0001, strains p<0.0001. N=5∼6 for both strains 3-7dpi. D: Immunofluorescent staining of PDGFRɑ (red), Ki67(green), and nuclei by DAPI (blue) of 5dpi muscle sections. Scale Bar: 200 µm. E: Quantification of proliferating FAPs on muscle section from 5dpi muscle. At 5dpi, FID muscles showed a reduction in Ki67^+^ FAPs comparing to *Igf1^fl/fl^* muscles. Unpaired t=test: p=0.0013. N=4 for both strains. F: Primary FAP cell culture phase-contrast images at the day of plating, 2-6 days post-plating (dpp). G: Quantification of cells per field 0-6 days post-plating shows the inability for cells from FID mice to proliferate. FAPs without IGF-I shows impairments in growth in culture. Two-way ANOVA: Interactions p<0.0001, timepoints p<0.0001, strains p<0.0001. N=6 for both strains. H: Immunofluorescent staining of primary FAPs with GFP (green), Ki67 (red), and DAPI (blue). Cells were exposed to rIGF-I, AEW541, or both for 24 hours Scale bar: 200µm. I: Quantification of proliferating FAPs in primary culture. Primary FAPs from FID muscles had lower Ki67^+^ cells compared to FAPs from *Igf1^fl/fl^* muscles. Exposure to rIGF-I increased proliferation in FAPs from *Igf1^fl/fl^* muscles, which was blocked by addition of AEW541. There was no effect of rIGF-I on proliferation of FAPs from FID muscles. Two-way ANOVA: Interactions ns, treatments p=0.009, strains p<0.0001. N=4 for both strains. J: Immunoblotting of total and phosphorylated Akt and ERK1/2 in primary FAPs exposed to rIGF-I, AEW541, or both for 30 minutes. GAPDH serves as loading control. Scale bar: 100µm or 200µm. An unpaired student t-test or one-way ANOVA followed by Dunnett’s multiple comparison test or two-way ANOVA (strains and injury time) followed by Šídák post-hoc test were used for statistical analysis. # represent comparison between timepoints and * represents comparison between strains. * or # p<0.05, ** or ## p<0.01, *** or ### p<0.001, **** or #### p<0.0001.

### Gene expression analysis reveals suppression of ECM production and cell proliferation in FAPs lacking IGF-I

To expand our understanding of the impact IGF-I has on FAP activity, we used Bulk-RNA Sequencing (BulkRNASeq) of primary FAPs isolated from regenerating muscles 3 dpi to capture differences between strains when the FAPs were most dynamic (Figure 4A). A total of 20,937 genes were identified, and with thresholds set to 1-fold changes with p<0.05, we detected 427 genes that were upregulated and 179 genes that were downregulated in FAPs without IGF-I compared to FAPs with IGF-I (Figure 4B). FAP purity was confirmed by the high expression level of FAP markers (*Pdgfra* and *Ly6a*) and the absence of expression of markers for immune cells, satellite cells, or myonuclei (Figure S3A). Gene ontology (GO) analysis revealed that the highest scores for upregulated genes were part of immune-related pathways, suggesting that FAPs from FID mice adopted an inflammatory phenotype (Figure 4C, 4D). Intriguingly, out of the 179 genes that were downregulated, the majority of them clustered into Extracellular Matrix (ECM) related groups (Figure 4C), which when examined by heatmaps displayed a global shutdown of ECM production in FAPs, including multiple collagen genes, fibronectin (*Fn1*), and laminin 2 (*Lama2*) (Figure 4E). In addition to reductions in ECM genes, downregulation of *Klf4* was evident in FID FAPs, which is a key regulator in fibrogenesis (Chandran *et al*, 2021) (Figure 4F). These changes were accompanied by a reduction in several genes promoting proliferation, consistent with the assessment of cell numbers during regeneration (Figure 4F). Because FAPs can adopt fibrogenic or adipogenic lineages (Joe *et al*., 2010; Uezumi *et al*., 2010), we examined the expression of genes associated with adipogenesis to determine if there was a shift if the fate of FAPs. Unlike the downregulation of ECM genes in FID FAPs, there was no consistent pattern in adipogenic gene expression, indicating an absence of trans-differentiation in lieu of a fibrogenic fate (Figure 4G). Validation of RNASeq data was achieved through qRT-PCR for genes representative of ECM, adipogenesis, cell proliferation, and immune responses (Figure S3B). Overall, the FAP gene expression profile in the absence of IGF-I exhibited a suppression of pathways needed to mediate muscle remodeling during regeneration.

**Figure 4.**
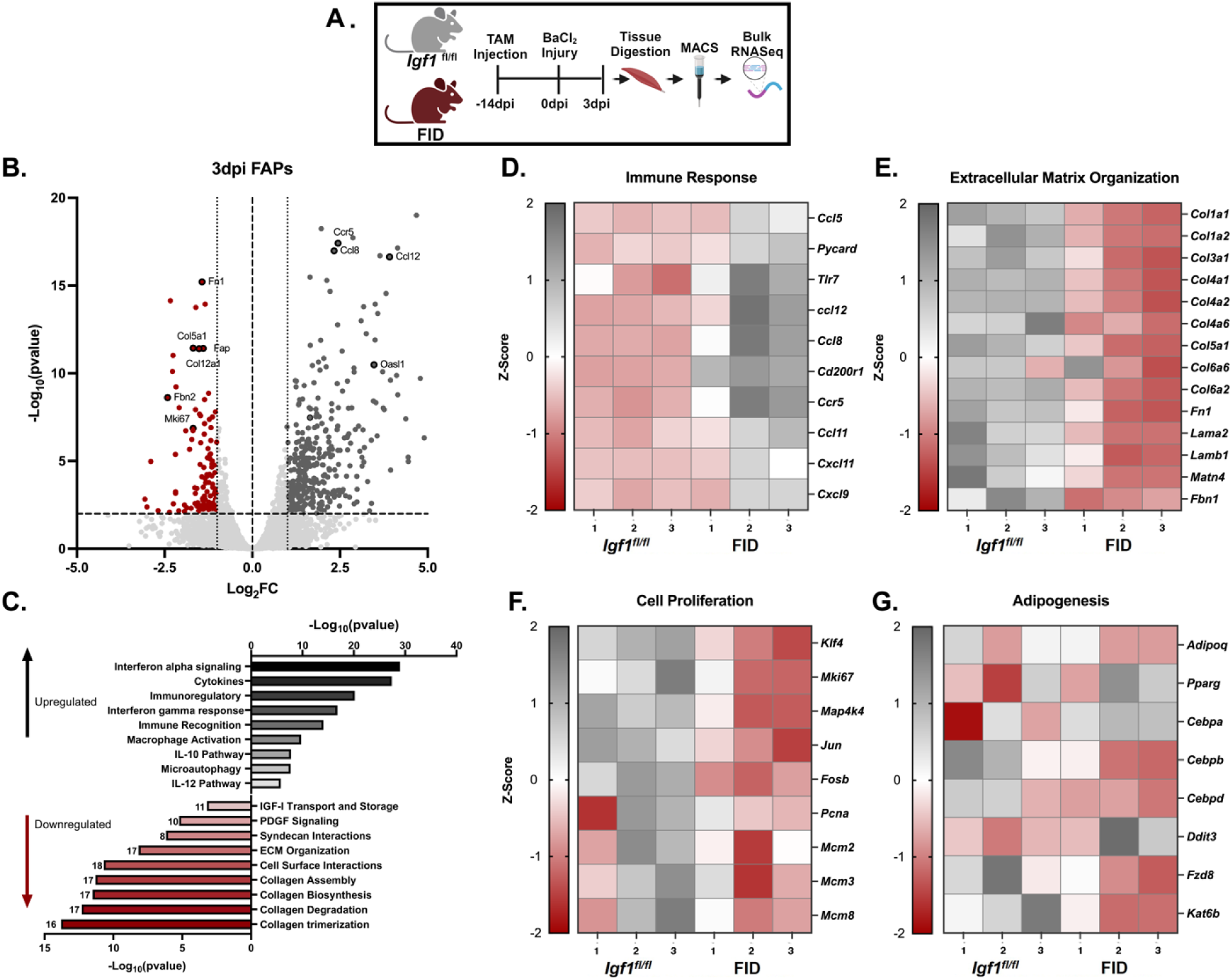
RNASeq of isolated FAPs reveals the necessity of IGF-I for FAP differentiation. A: Schematic illustration of Bulk RNA sequencing. Primary FAPs are isolated from 3dpi TA using MACS sorting. After isolating primary FAPs, RNA was isolated and downstream sequencing was performed. B: Volcano plot of all the genes that were captured in RNASeq. Log_2_ Fold change of 1 and p-value<0.05 were used as cutoffs. Upregulated genes are shown in dark grey and downregulated genes are shown in dark red. C: Pathway analysis of the primary FAPs isolated from 3dpi TA muscle using MACS. D-G: Heatmaps of differential gene expression for immune response, ECM organization, cell proliferation, and adipogenesis. Z-score ranges from −2 to 2. Downregulation showed in dark red, and upregulation showed in grey.

### Loss of FAP IGF-I suppresses ECM deposition and fibroblast differentiation

During muscle regeneration, FAPs deposit a significant amount of ECM that serves as a scaffold for the muscle as damaged fibers are degraded and new ones are formed (Kotsaris *et al*, 2023; Lemos *et al*., 2015). In models where FAPs are targeted, ECM content is reduced (Hammers, 2022; Kotsaris *et al*., 2023; Lemos *et al*., 2015). Prompted by the RNASeq profile of FAPs from FID mice, we assessed the effects of IGF-I loss from FAPs on ECM content during regeneration by Picro-Sirius-Red (PSR) staining (Figure 5A). The robust transient increase in ECM content found in regenerating *Igf1^fl/fl^* muscles did not occur in FID muscles (Figure 5B, C) in concurrence with the suppressed ECM gene expression (Figure 4E). However, this observation could simply be due to the reduced presence of FAPs in regenerating FID muscles rather than any alteration in FAP differentiation into fibroblasts/myofibroblasts (Figure 3B, C). To determine if fibroblast differentiation was directly impacted by loss of IGF-I from FAPs, we stimulated primary FAPs with TGF-β (5 ng/mL) to induce fibroblast differentiation (Contreras *et al*., 2019) (Figure 5D). Experiments were initiated at the same cell confluency to control for the effects of FAP IGF-I on cell proliferation. FAPs from *Igf1^fl/fl^* muscles adopted a fibroblast-like morphology 12 hours after TGF-β treatment, whereas those from FID muscles were delayed by 24 hours in the appearance of fibroblast morphology (Figure 5E). Expression analysis of FAPs for ECM genes revealed comparable upregulation of collagens (*Col1a4*, *Col4a1*) 24 hours post TGF-β induction indicated that the differentiation pathway was intact, yet expression of fibronectin (*Fn1*) remained low in FAPs from FID muscles (Figure 5F-H) (Jones *et al*, 2019). To further understand the nature of the delay in FID FAPs, immunoblotting for TGF-β signaling and markers for FAPs, myofibroblasts, and adipocytes were performed (Figure 5J). TGF-β mediated phosphorylation of Smad2 was similarly elevated in FAPs from both FID and control *Igf1^fl/fl^* mice (Figure 5J, S4H). However, there was a delay in the production of alpha smooth muscle actin (αSMA), a marker for myofibroblasts, in FID FAPs until 12 hrs post-induction (Figure 5J,K). Further, TGF-β incubation caused an immediate reduction in PDGFRα indicative of FAPs entering the fibroblast lineage (Kotsaris *et al*., 2023), yet there was a retention of PDGFRα in FID FAPs throughout the 48 hr TGF-β incubation (Figure 5J, L). This suggests that FID FAPs were resistant to differentiation, remaining in the PDGFRα^+^ progenitor state. Finally, there was no evidence of perilipin accumulation in any FAPs (Figure 5J), supporting the absence of endogenous pro-adipogenic transdifferentiation in FID FAPs. In sum, the FAPs from FID mice were responsive to strong fibrogenic stimuli, but they were delayed in their differentiation, supporting the observations *in vivo* of regeneration deficits.

**Figure 5.**
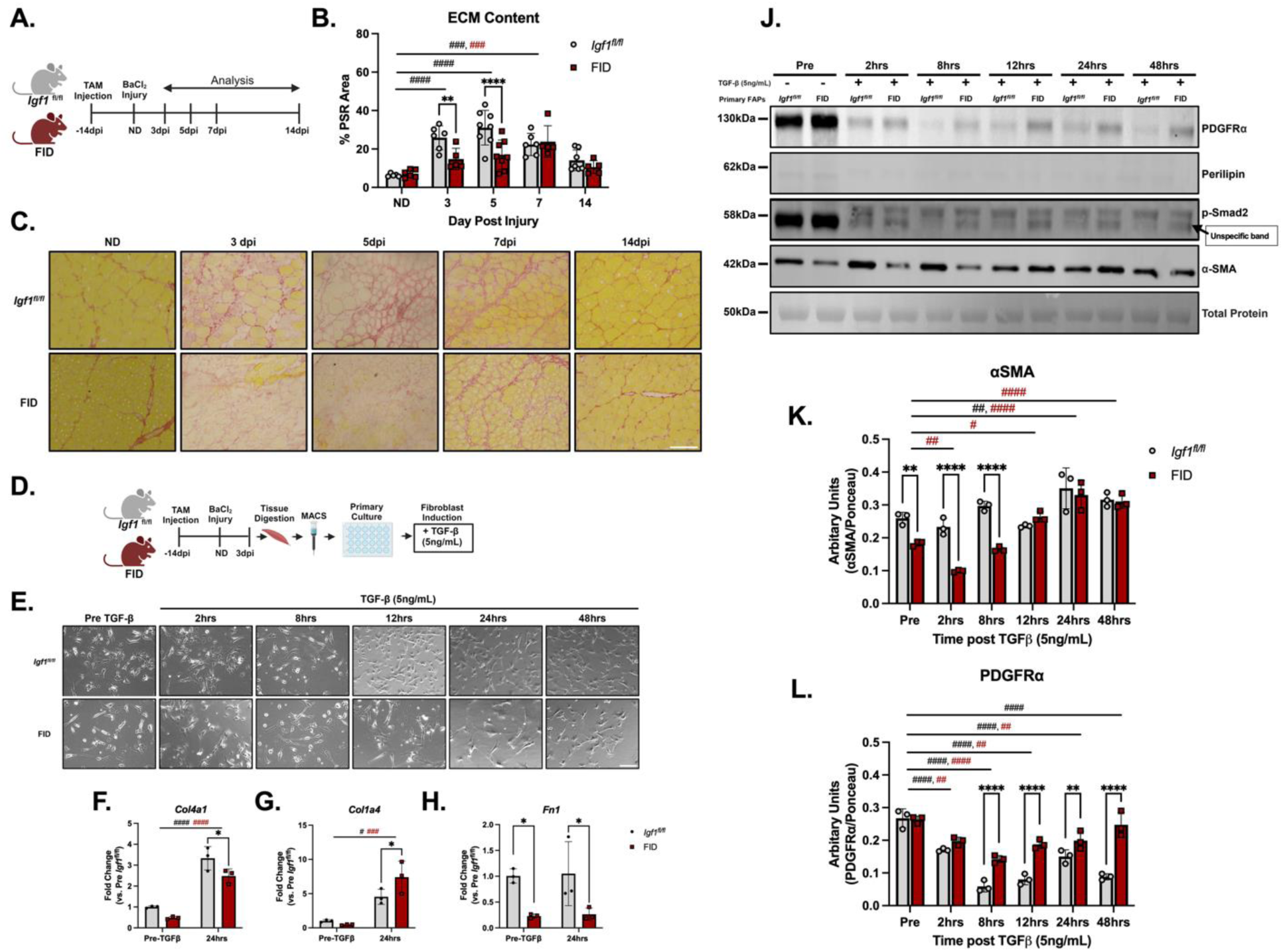
IGF-I from FAPs is required for ECM deposition during regeneration *in vivo* and fibroblast differentiation *in vitro*. A: Graphical illustration of the experimental design for muscle regeneration and time of analysis. B: Quantitative analysis of the extracellular matrix content during muscle regeneration by area positive for picro-sirius red (PSR) staining. At 3 and 5 dpi, FID muscles exhibit less ECM content compared to regenerating *Igf1^fl/fl^*muscles. Only *Igf1^fl/fl^* muscles showed a significant increase in ECM content at 3 and 5 dpi. Muscles from both strains exhibited elevated ECM at 7 dpi. Two-way ANOVA analysis: interaction p=0.0062, timepoint p<0.0001, strains p=0.0019. N= 5∼8 for each strain at all timepoints. C: Picro-Sirius-Red (PSR) staining of TA muscles from ND to 14 dpi. Scale bar: 200µm. D: Graphical illustration of primary FAP isolation for TGF-β (5ng/mL) induced fibroblast differentiation. E: Phase-contrast images of primary FAPs at desired confluency (∼60%) prior to TGF-β induction, and at multiple timepoints during the following 48 hrs post TGF-β induction. F: qRT-PCR analysis of *Col4a1* in FAPs from pre and 24 hrs post incubation. Two-way ANOVA analysis: interaction p=ns, timepoint p<0.0001, strain p=0.0069. N=3 for both strains at both timepoints. G: qRT-PCR analysis of *Col1a4* in FAPs from pre and 24 hrs post incubation. Two-way ANOVA analysis: interaction p=ns, timepoint p<0.0001, strain p=0.1648. N=3 for both strains at both timepoints. H: qRT-PCR analysis of *Fn1* in FAPs from pre and 24 hours post incubation. Two-way ANOVA analysis: interaction p=ns, timepoint p=ns, strain p=0.0031. N=3 for both strains at both timepoints. J: Immunoblotting analysis of protein isolated from primary FAPs following TGF-β treatment. Total protein quantification was obtained from ponceau staining. K: Quantification of αSMA obtained from immunoblotting analysis. Two-way ANOVA analysis: interactions p<0.0001, timepoints p<0.0001, strains p<0.0001. For *Igf1^fl/fl^*, p=0.0041 for comparison between pre-incubation and 24hrs post-incubation. For FID, p<0.05 or p<0.001 for comparison between pre-incubation and 12hrs, 24hrs, and 48hrs post-incubation. N=3 for each timepoints for both strains. L: Quantification of PDGFRα obtained from immunoblotting analysis. Two-way ANOVA analysis: interactions p<0.0001, timepoints p<0.0001, strains p<0.0001. p<0.01 or p<0.0001 at 8,12,24 and 48hrs between strains. *Igf1^fl/fl^* with p<0.0001 between all timepoints and pre-incubation within. FID with p<0.01 or p<0.0001 between all timepoints and pre-incubation expect 48hrs. N=3 for each timepoints for both strains. Scale bars: 200µm. Two-way ANOVA (strains and timepoints) followed by Šídák post-hoc test used for statistical analysis. #: comparison between timepoints within strain and *: comparison between strains, with red # indicating comparisons for FID, and black # indicating comparisons for *Igf1^fl/fl^*. * or # p<0.05, ** or ## p<0.01, *** or ### p<0.001, **** or #### p<0.0001.

### Loss of FAP IGF-I reduces adipocyte deposition and adipogenic differentiation

In addition to being the source of ECM deposition and muscle fibrosis, FAPs are also the primary progenitor cells for intramuscular adipocytes (Kopinke *et al*., 2017; Uezumi *et al*., 2010). Because the FAPs from FID mice had a suppressed fibrogenic response in regenerating muscle, a similar blunted response to adipogenic stimuli might also occur. To determine if IGF-I from FAPs altered intermuscular fat formation, an injection of 50% glycerol was performed, which induces adipocyte formation (Johnson *et al*, 2022; Kopinke *et al*., 2017; Norris *et al*., 2023; Norris *et al*, 2024) (Figure 6A). Only female mice were utilized for this evaluation, due to the greater adipogenic response in C57BL/6 female mice (Johnson *et al*., 2022; Kopinke *et al*., 2017; Norris *et al*., 2023). Formation of intermuscular adipocytes was evident by 7 dpi, indicated by immunostaining for perilipin (Figure 6B, C). By 21 dpi, accumulation of fat was more extensive in *Igf1^fl/fl^*muscles than in FID muscles (Figure 6B, C). Unlike BaCl_2_ induced injury, there were no differences in the masses of muscles between strains as glycerol injury resolved (Figure S4A, B). Taken together, loss of IGF-I from FAPs resulted in a suppression of adipocyte accumulation following glycerol induced injury without altering the myogenic aspects of damage resolution.

**Figure 6.**
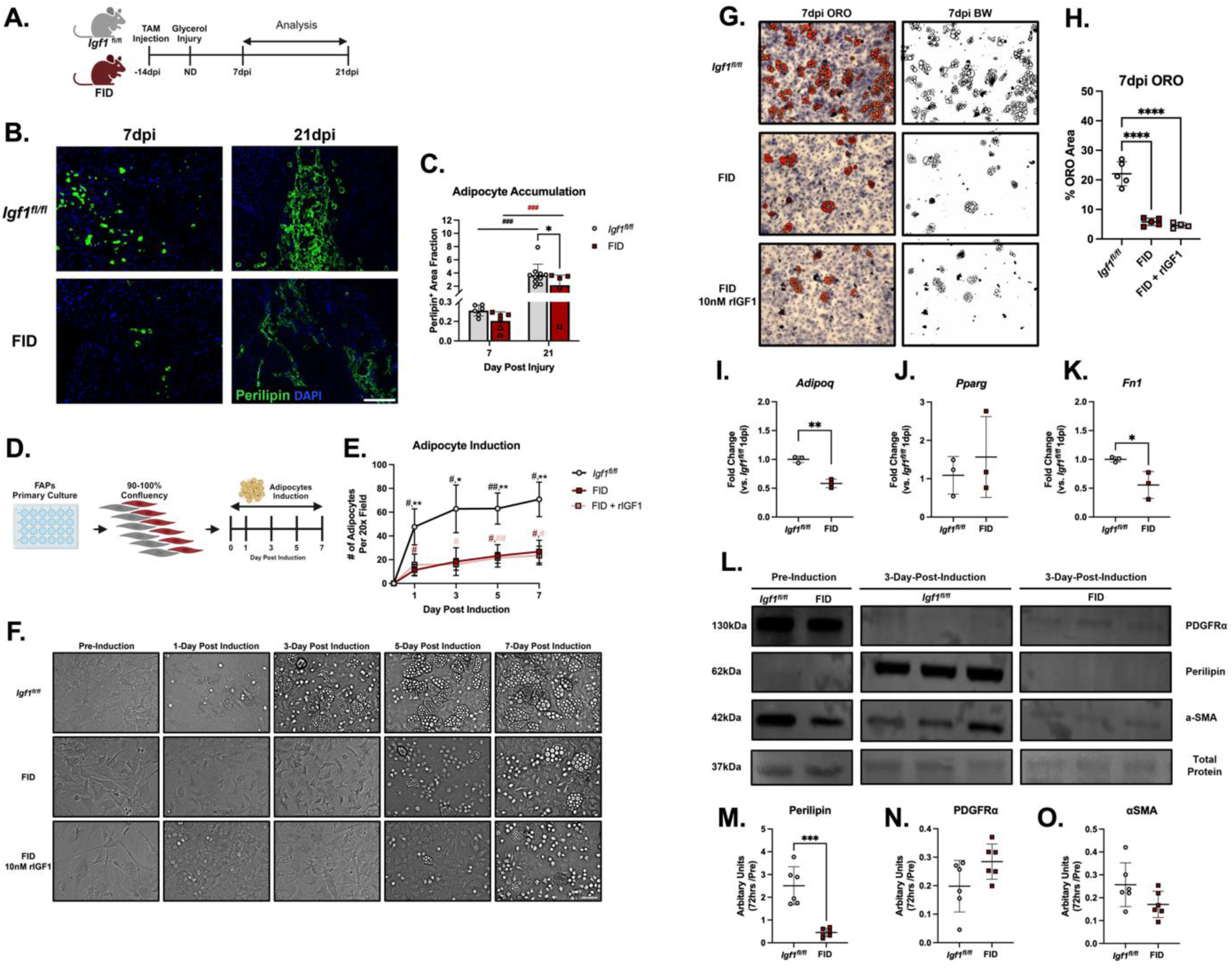
Impact of IGF-I on adipocyte accumulation post glycerol injury and in-vitro adipocyte differentiation. A: Schematic illustration of the glycerol injury model. B: Immunofluorescent staining of Perilipin (green) and nuclei (DAPI, blue) on cryosections of skeletal muscle at 7 and 21 dpi. Scale Bar: 200µm. C: Quantification of adipocytes at 7 and 21 dpi. FID muscles had significantly lower adipocyte accumulation at 21 dpi. Two-way ANOVA analysis: interaction p=ns, timepoints p<0.0001, strains p=0.0992. N=6∼10 muscles for both strains at 7 and 21 dpi. D: Graphical illustration of the adipocyte induction experimental design in-vitro. E: Quantification of the number of adipocytes after induction. Robust increases in adipocyte formation in cultures from *Igf1^fl/fl^* muscles was significantly blunted in primary FAPs from FID muscles, regardless of IGF-I treatment. Two-way ANOVA analysis: interactions p<0.0001, timepoints p<0.0001, strains p<0.0001. *Igf1^fl/fl^* p<0.0001 between all timepoints post induction and 0dpi. N=4∼5 for all timepoints and strain conditions. F: Phase contrast or live cell imaging (GFP) of primary FAPs before and following adipocytes induction. Scale Bar: 200µm. G: Oil-Red-O staining at 7-day post induction. Right column shows black and white 8-bit high contrast image for better visualization. H: Quantification of the ORO+ area at 7dpi with FAPs from control, FID, and FID with supplemental IGF-I. FAPs from FID muscles showed a significant reduction if ORO+ area that was not restored with exogenous recombinant IGF-I. One-way ANOVA analysis p<0.0001. N=4 per strain per condition. I - K: qRT-PCR analysis of FAPs 24hrs post induction. Unpaired t-test are performed for *Adipoq*, *Pparg*, and *Fn1*. *Adipoq* p=0.0013, *Pparg* p=0.5175, *Fn1* p=0.032. N=3 for both strains. L: Immunoblotting of primary FAPs at 3dpi probing for PDGFRα, Perilipin, and αSMA. Ponceau was used a total protein loading control. M-O: quantification of the immunoblotting of primary FAPs at 3dpi. Unpaired t-tests performed for Perilipin, PDGFRα, and αSMA. Perilipin p=0.0002, PDGFRα p=0.0824, αSMA p=0.0868. N=6 for both strains. Scale bars: 200µm. An unpaired student t-test or one-way ANOVA followed by Tukey test or two-way ANOVA (strains and injury time) followed by Šídák post-hoc test were used for statistical analysis. #: comparison between timepoints within strain and *: comparison between strains, with red # indicating comparisons for FID, and black # indicating comparisons for *Igf1^fl/fl^*. * or # p<0.05, ** or ## p<0.01, *** or ### p<0.001, **** or #### p<0.0001

To more closely examine the dependence of FAP IGF-I on adipogenesis, primary FAPs isolated from muscles of both strains were plated and grown until optimal confluency (95-100%), then treated with adipogenic induction media (0.25µM dexamethasone, 0.5mM isobutyl-methylxanthine, 5ug/mL Insulin and 5µM troglitazone) (Figure 6D). Lipid droplets began to accumulate in FAPs from *Igf1^fl/fl^* mice by 1-day post-induction in contrast to FAPs isolated from FID muscles where lipid droplet accumulation was not apparent until 5 days post-induction (Figure 6E, F). By 7 days in induction media, ∼70% of the cells from *Igf1^fl/fl^* mice adopted adipocyte morphology with positive oil red O (ORO) staining, whereas far fewer cells in FID cultures were positive for ORO (Figure 6E-H).

IGF-I has been used to induce adipogenesis in mesenchymal stem cells (Scavo *et al*, 2004; Smith *et al*, 1988; Wabitsch *et al*, 1995). To determine if exogenous IGF-I could rescue adipogenic differentiation in FID cultures, 10nM rIGF-I was added to the induction media in replacement of insulin. This failed to alter the time course of adipogenic differentiation in FAPs from FID muscles (Figure 6E-H). Since *Igf1r* expression is highest during early stages of adipogenesis and begins to downregulate after 3dpi (Scavo *et al*., 2004; Smith *et al*., 1988; Zizola *et al*, 2002), we investigated biochemical changes within that window. Expression of adiponectin (*Adipoq*) was lower in FAPs from FID muscles 24 hours after adipogenic induction, consistent with suppressed adipogenesis. Even so, there were no differences in expression of *Pparg*, a master regulator of adipocyte differentiation (Chawla *et al*, 1994), between FAPs from FID or *Igf1^fl/fl^* muscles (Figure 6J), suggesting that IGF-I regulates adipogenesis independently of PPARG actions. *Fn1* expression served as a marker of fibrogenic transdifferentiation (Figure 6K), and it remained lower in FID cultures similar to the findings in fibrogenic media.

Markers of differentiation were quantified by immunoblotting of cell lysates 3 days post induction (Figure 6L-O). A robust increase in perilipin levels indicative of adipogenic differentiation was observed in FAPs from *Igf1^fl/fl^*muscles, but perilipin remained low in FAPs from FID muscles (Figure 6L, M). Pre-induction cells exhibited strong PDGFRα protein levels, regardless of strain, and levels decreased after induction but to a lesser extent in FID cells (Figure 6L,N), suggesting that FID FAPs were more likely to remain in the PDGFRα^+^ progenitor state. The myofibroblast marker, αSMA, was strong in cells from both strains prior to induction, with 75-85% reduction after adipogenic induction (Figure 6L,O). Thus, similar to the response of FID FAPs to fibrogenic stimuli, they exhibited a poor response to adipogenic stimuli and did not transdifferentiate into other fates. Overall, IGF-I from FAPs is important for adipocyte formation after glycerol induced muscle injury and for *in vitro* adipogenesis.

### Single cell expression analysis reveals loss of fibrotic FAPs and adoption of senescent phenotype in homeostatic FID muscles

Our assessment of IGF-I loss from FAPs examined dynamic processes and showed that without an autocrine source of IGF-I, FAPs are unable to properly differentiate or participate in muscle remodeling. However, it was not clear if they entered this state upon a stimulus or if they had already adopted this state after IGF-I reduction. To probe the effects of FAP *Igf1* deletion at the cellular level in homeostasis, single-cell RNA sequencing (scRNAseq) was performed on TA muscles pooled from each strain 21 days after TAM induction (N=5 male mice per strain). Because muscle fibers from FID mice displayed minimal changes in their properties without damage (Figure S1), scRNAseq afforded a closer examination of other cell types in the muscle tissue. Approximately 15,000 cells were assessed per strain with 40,000 reads per cell. After UMAP clustering, 9 cell populations were identified in homeostatic muscle according to their unique cell markers (Figure 7A, Figure S9A, S9D-E) (Oprescu *et al*, 2020b). FID samples displayed a notable reduction in the proportion of FAPs, fibroblasts, and monocytes, and a proportional increase in endothelial cells (Figure 7B). FAPs, fibroblasts, and monocytes were the predominant sources of *Igf1* expression (Figure 7C), as previously established (Kok & Barton, 2021). Because deletion of exon 4 did not alter transcription levels of *Igf1* (Figure 1G, S3B), the expression signature did not reveal which cells lost exon 4. There were no obvious compensatory changes in *Igf1* expression in other cell types in FID samples (Figure 7C), suggesting that in homeostatic muscle, loss of FAP IGF-I does not trigger increased expression from other cells.

**Figure 7.**
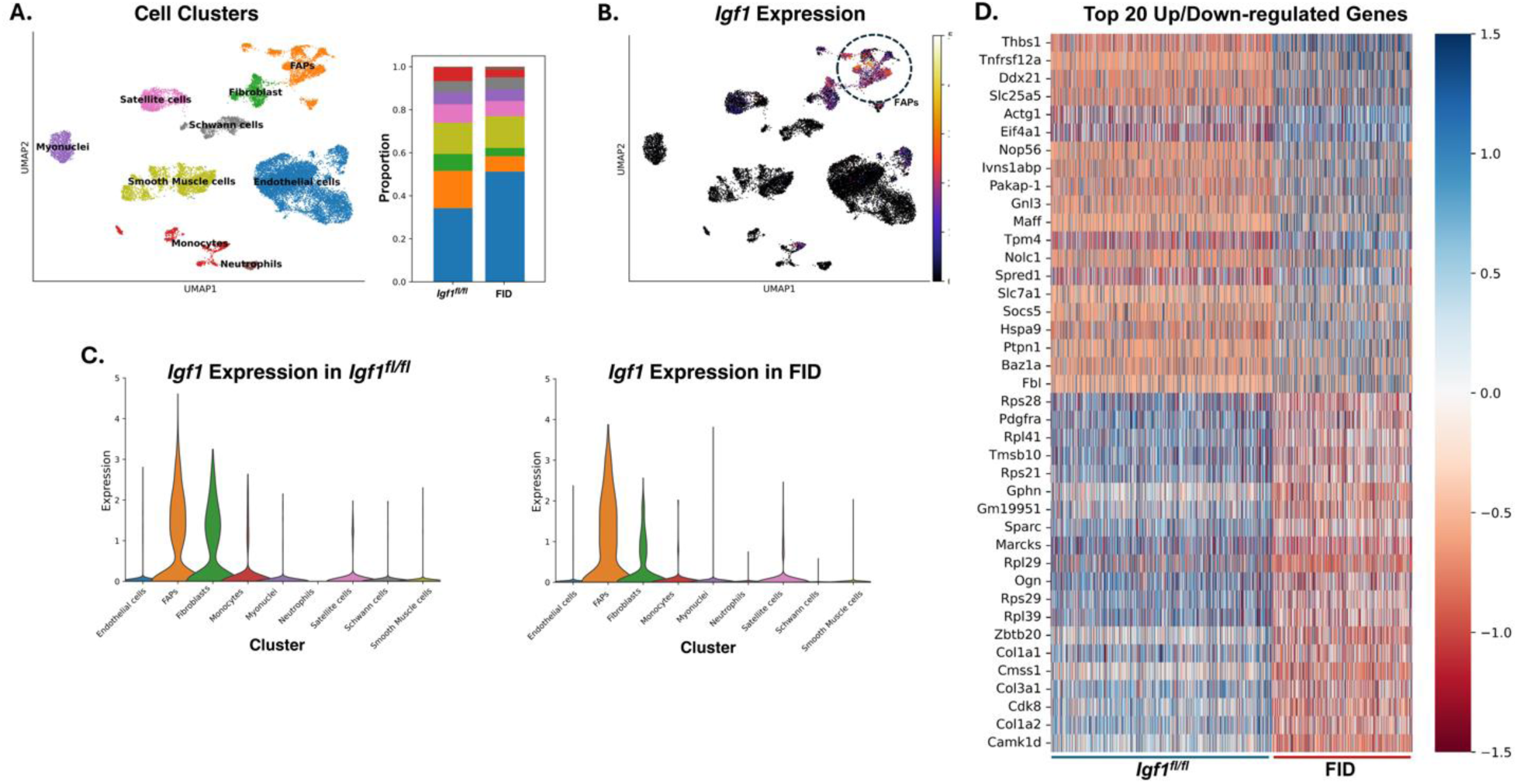
scRNASeq analysis of homeostatic muscles. A: Cell clusters identified by gene markers (details in Figure sXX). FAPs and fibroblasts are less abundant in FID muscle samples compared to cell proportions in *Igf1^fl/fl^* muscle samples for after clusters are defined. B: *Igf1* expression is detected in FAPs, fibroblasts, and monocytes. C: Violin plots of Igf1 expression across clusters show the majority of *Igf1* expression occurs in FAPs and fibroblasts, and that there is no compensaorry change in expression in other cell clusters when *Igf1* is ablated from FAPs in FID mice. C: Comparison of the top 20 up- and down-regulated genes in FAPs.

To further understand the effects of *Igf1* deletion in FAPs, downstream analysis was performed on the FAP cluster. We observed a significant and global reduction of collagen gene expression in FID FAPs, supporting that ECM production was already altered in homeostatic muscle (Figure 7D). Examination of the top 20 up-and down-regulated genes also revealed the striking finding is the elevation of *Thbs1* and *p53*. *Thbs1* has been shown to have tumor suppression effects by stabilizing p53, which drives cell cycle arrest and senescence (Lawler *et al*, 2001; Linderholm *et al*, 2004; Skeen *et al*, 2006; Sundaram *et al*, 2011). Downregulation of *Cdk8*, *Rps28*, *Rps29*, and *Rpl29*, crucial genes for cell proliferation or ribosome-dependent protein translation (Kim *et al*, 2019; Kirn-Safran *et al*, 2007; Mirabello *et al*, 2014; Smith & Eppig, 2009; Szilagyi & Gustafsson, 2013) was also evident. Examination of the FAP cluster revealed four sub-populations based on unique gene signatures associated with their actions. These included (1) fibrotic FAPs with the highest collagen gene expression, (2) progenitor FAPs are expressing *Dpp IV,* (3) activated FAPs with high transcriptional activity of *Sox6*, and (4) microenvironmental FAPs with expression of critical ECM genes (Babaeijandaghi *et al*, 2023; Krebsbach & Villa-Diaz, 2017; Mann *et al*, 2011) (Figure 8A-B, Figure S10A-B). Fibrotic FAPs were the most affected both in number and gene expression, consistent with the general reduction of collagen gene expression in the total FAP pool (Figures 4C-D; 7D). Next, we performed gene ontology and pathway analysis to gain insight on the impact IGF-I may have on biological processes in FAPs (Kanehisa *et al*, 2017). Similar pathways emerged, with ECM structure, p53 regulation, and protein translation most evident (Figure 8C-E). Taken together, the loss of FAP IGF-I led to a shift in the FAP expression profile toward an inactive, possibly senescent, state. This shift may underly the inability for FID FAPs to participate in muscle regeneration.

**Figure 8.**
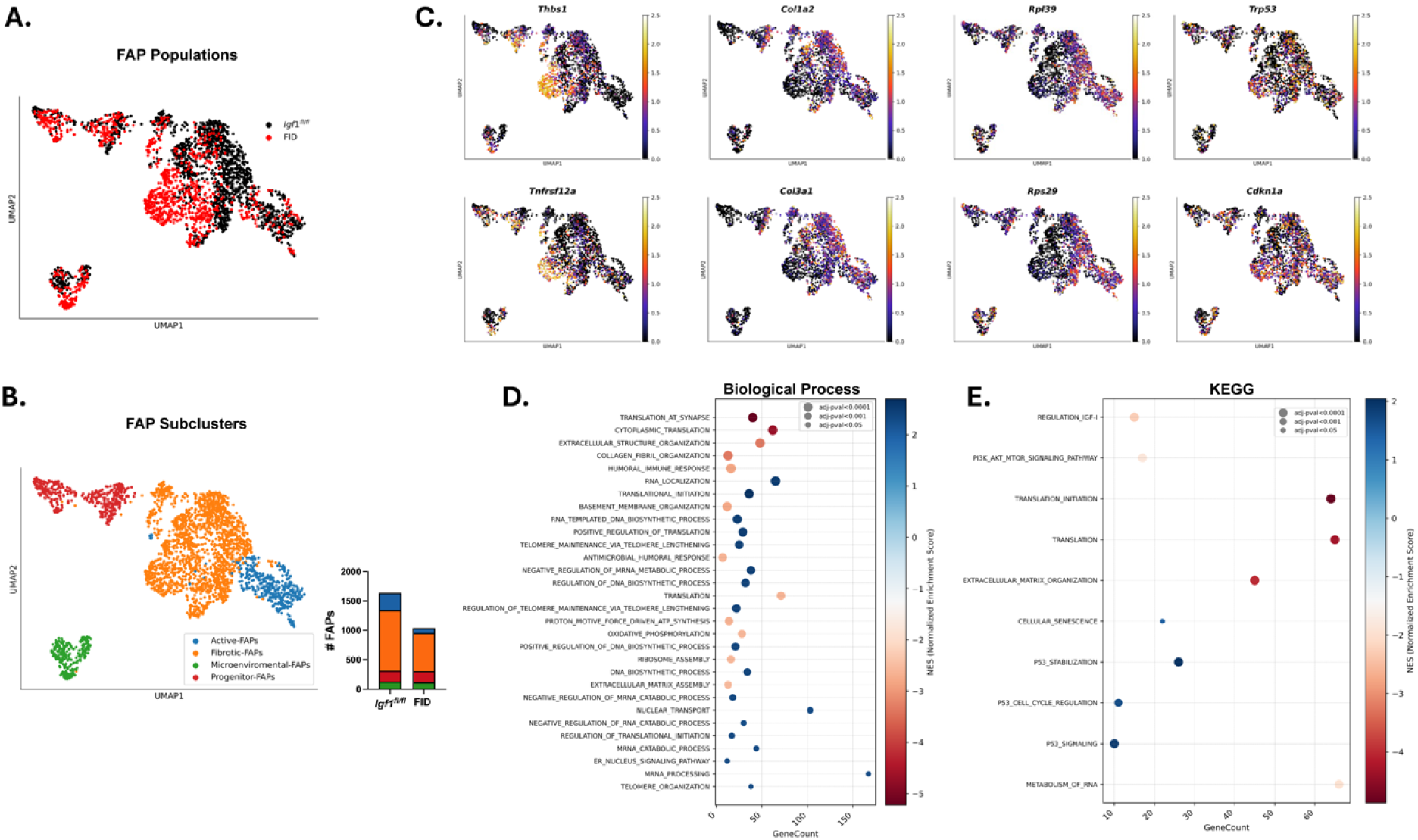
Single cell gene expression analysis of FAPs. A: Overlay of FAPs from FID and *Igf1^fl/fl^* muscle reveal a separation in the profiles of fibrotic FAPs from FID samples away from those from *Igf1^fl/fl^*muscle. B: Identification of four FAP sub-populations: (1) fibrotic FAPs with the highest collagen gene expression, (2) progenitor FAPs expressing *Dpp IV,* (3) activated FAPs with high *Sox6* expression, and (4) microenvironmental FAPs with ECM gene expression. There are ∼30% fewer FAPs in FID muscles, with active and fibrotic FAPs being the most affected. C: UMAPs for representative most changed genes. The two highest gene expression changes (*Thbs1* and *Tnfrsf12a*) are most pronounced in the fibrotic FAPs of FID muscles. The lowest expressing collagen genes (*Col1a2* and *Col3a1*) are distributed throughout the cluster, as are the low expression of the protein synthesis genes (*Rpl39* and *Rps29*). High expression of *Trp53* is also distributed throughout the cluster. D: Biological process analysis in FAP differential expression displays reduced translation as well as ECM and basement membrane formation in FID FAPs. Increases in pathways associated with telomere maintenance as well as regulation of translation are also evident. F: KEGG analysis reveals the FAPs from FID muscles have suppression of translation, ECM organization, RNA metabolism and IGF-I pathways, and enhancement of p53 and cellular senescence.

### IGF-I ablation induces a senescent-like state in fibro-adipogenic progenitors

Cellular senescence in FAPs has been previously observed during skeletal muscle regeneration and under pathological conditions (Moiseeva *et al*, 2023; Saito *et al*, 2020). Here, we found that FID FAPs exhibit transcriptional profiles closely resembling those of senescent FAPs (Moiseeva *et al*., 2023) marked by elevated inflammatory signaling, reduced expression of cell proliferation markers, matching a senescence cell phenotype (Figures 4D; 8; S3C-D). These findings prompted us to examine senescence in FAPs more closely. In homeostatic muscle, 22% of FID FAPs displayed co-localization of PDGFRα with β-Galactosidase (β-Gal) immunofluorescence compared to 5% of the FAPs in WT muscles, indicating a pre-existing senescent phenotype in FID FAPs independent of acute muscle injury (Figure 9A-C). Following BaCl_2_ injury, more than 60% of FAPs lacking IGF-I were senescent at 7 dpi, compared to only 20% of *Igf1^fl/fl^* FAPs (Figure 9A-C), with little amplification of FAP numbers in regenerating FID muscles. By 28 dpi, FAP numbers returned to homeostatic levels in both strains. However, the proportion of senescent FAPs persisted above 60% in FID muscles. While cellular senescence is typically observed as a physiological event during muscle regeneration (Moiseeva *et al*., 2023), the absence of IGF-I appears to predispose FAPs to a senescent state prior to injury. Further, the population of senescent FAPs may grow following regeneration in FID muscles, potentially having a negative effect on resolution of damage.

**Figure 9.**
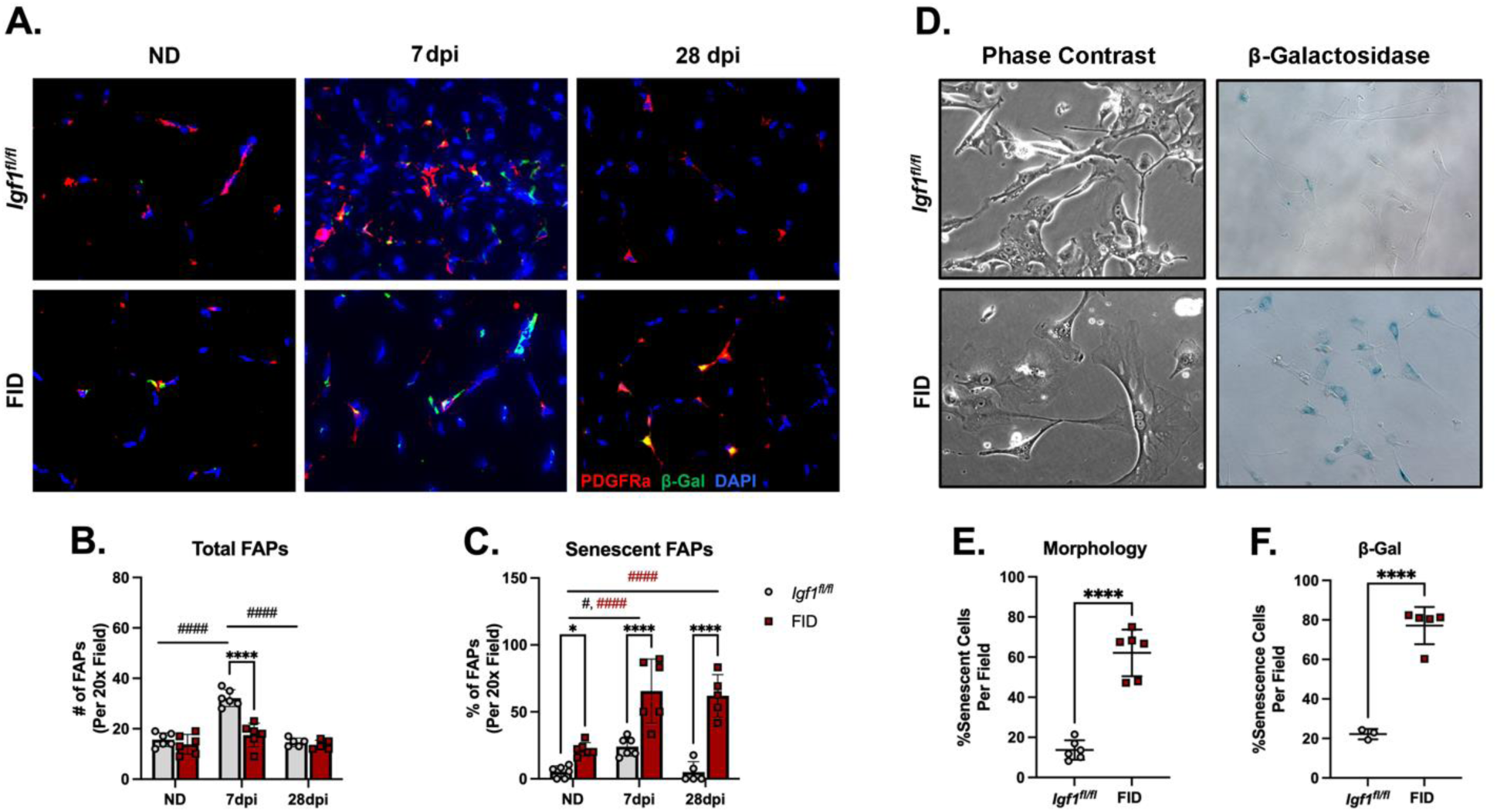
IGF-I ablation induces cellular senescence in Fibro-Adipogenic Progenitors A: Immunofluorescent staining of PDGFRα (red), β-galactosidase (green), and DAPI (blue) on cryosections of skeletal muscle from ND and 7dpi. Scale Bar: 100µm. B: Quantification of FAP numbers show that the transient expansion of FAPs in regenerating *Igf1^fl/fl^* muscles is not apparent in FID muscles.Two-way ANOVA analysis: interaction p<0.0001, timepoints p<0.0001, strains p<0.0001. N=6 for both strains at all timepoints. C: Quantification of β-galactosidase positive FAPs as an index of senescence show that FAPs from FID mice have higher index of senescence in homeostatic and regenerating muscles. Two-way ANOVA analysis: interaction p=0.0299, timepoints p<0.0001, strains p<0.0001. N=6 for both strains at all timepoints. D: Brightfield image of plated primary FAPs, and after β-Gal staining. E: Quantification of the proportion of senescent FAPs based on their morphology show significantly higher senescence in FAPs from FID mice. Unpaired t-test: p<0.0001. N=6 for both strains. F: Quantification of the proportion of β-Gal positive FAPs based on their blue color development show significantly higher senescence in FAPs from FID mice. Unpaired t-test: p<0.0001. N=4 for *Igf1^fl/fl^*, N=5 for FID. An unpaired student t-test or two-way ANOVA (strains and injury time) followed by Šídák post-hoc tests were used for statistical analysis. #: comparison between timepoints within strain and *: comparison between strains, with red # indicating comparisons for FID, and black # indicating comparisons for *Igf1^fl/fl^*. * or # p<0.05, ** or ## p<0.01, *** or ### p<0.001, **** or #### p<0.0001

To further evaluate this senescent phenotype in FAPs lacking IGF-I, we isolated primary FAPs from both mouse strains. FID FAPs showed morphological characteristics typical of senescent cells, including an enlarged cytoplasm, a flattened cell shape, and multinucleation (Beck *et al*, 2020; Ben-Porath & Weinberg, 2004) (Figure 9D,E). Finally, we measured β-Gal activity in primary FAPs: while *Igf1^fl/fl^*FAPs exhibited minimal β-Gal activity, most FAPs lacking IGF-I were positive for β-Gal activity (Figure 9D,F). Taken together, FAPs from FID muscles adopt senescent properties *in vitro* and *in vivo* in homeostatic conditions, and this is exacerbated during muscle regeneration.

## Discussion

Research in the past decade has established that FAPs play a critical role in muscle homeostasis, regeneration, aging, and even neuromuscular junction integrity (Joe *et al*., 2010; Kim *et al*., 2022; Lukjanenko *et al*, 2019; Uezumi *et al*., 2021; Wosczyna *et al*., 2019). It is also well known that FAPs secrete IGF-I, which is a well-documented factor in contributing to skeletal muscle biology (Barton-Davis *et al*., 1998; Kok & Barton, 2021; Luo *et al*., 2024; Musaro *et al*., 2001; Vassilakos *et al*., 2019). Both the sources and timing of IGF-I production are central to skeletal muscle growth and regeneration. This motivated the central questions of this study, to determine if the FAP IGF-I source was needed for efficient muscle regeneration in general, and to determine if FAP actions were affected in particular. Using PDGFRα specific inducible Cre recombinase expression in mice enabled the virtually complete ablation of *Igf1* expression in FAPs. This resulted in defects in the early and late phases of regeneration, including smaller fibers, fewer satellite cells, and persistence of macrophages, implicating that IGF-I from FAPs altered the intercellular interaction during regeneration. While this could be a direct paracrine action of IGF-I on the multiple cell populations needed to resolved muscle damage, our examination of the FAPs themselves lead to a more likely scenario, in which IGF-I from FAPs is a critical autocrine factor, important for maintaining FAPs in homeostasis and during regeneration. Thus, we assert that FAP IGF-I indirectly affects the process of regeneration via its direct impact on FAPs. Multiple lines of evidence support these autocrine effects, including the lack of FAP expansion and diminished ECM deposition after BaCl_2_ injury, as well as reduced fat deposition after glycerol injury in muscles. Further, isolated FAPs exhibit extremely slow proliferation, blunted responses to fibrogenic and adipogenic media, and phenotypic hallmarks of senescence. Hence, the ablation of IGF-I from FAPs have essentially removed them from effectively participating in the regeneration process, mirroring the phenotype associated with the complete ablation of FAPs (Wosczyna *et al*., 2019).

With respect to the autocrine regulation of FAPs, several other pathways have been identified that alter FAP fate. Odd-skipped related transcription factor 1 (Osr1) is an important regulator of FAP activity during muscle regeneration. Loss of *Osr1* impedes FAP proliferation, enhances FAP apoptosis, impairs immune interactions, and upregulates extracellular matrix (ECM)-related genes (Kotsaris *et al*., 2023). We have delineated a contrasting phenotype with reduced extracellular matrix (ECM) production and enhanced immune pathway activity, without observable changes in *Osr1* expression. This suggests an independent pathway, potentially acting in opposition to Osr1-regulated mechanisms. Although IGF-I is not a transcription factor, it modulates the activity of multiple transcriptional regulators, such as Klf4 and ERK (Basisty *et al*, 2020; Chandran *et al*., 2021; Smith *et al*., 1988; Tosca *et al*, 2008; Yao *et al*, 2003; Zou *et al*, 2019). Our observed phenotype indicates the presence of a key regulator that governs global ECM production, warranting further investigation to elucidate its role and interaction with IGF-I signaling.

Upon activation following acute injury, FAPs develop primary cilia, a crucial event for FAP function in muscle tissue, as cilia-deficient FAPs exhibit resistance to adipogenic differentiation, in part, through the disruption of Hedgehog signaling (Kopinke *et al*., 2017). The expression of two primary cilia formation genes, *Arl13b* and *Ift88*, remained unaffected in FID FAPs, suggesting that IGF-I does not alter primary cilia. Nevertheless, IGF-I receptor (IGF1R) is localized on the primary cilia of mesenchymal stem cells (Dalbay *et al*, 2015), and it remains unclear if signaling crosstalk between IGF-I and other pathways is disrupted, such as Hedgehog signaling, enriched in primary cilia. Because we observed diminished regenerative capacity in FID muscles, rather than an enhanced myogenesis phenotype previously observed in the absence of primary cilia (Kopinke *et al*., 2017), it suggests that primary cilia formation is intact, or that the impact of IGF-I loss from FAPs overrides the potential benefit of primary cilia loss.

A major surprise in our study was the adoption of senescent hallmarks by the FAPs in homeostatic and regenerating FID muscles that was replicated in primary cultures of FAPs isolated from FID muscles. While a transient increase of senescent cells has been documented in muscles regenerating after acute injury, increases in the abundance of senescent cells are apparent in aging and diseased muscles, and impair efficient regeneration (Moiseeva *et al*., 2023). Senescence can occur in any of the cell types involved in regeneration; however, in our model, few, if any, cells other than FAPs displayed many hallmarks of senescence. A major contribution to the delay of muscle regeneration by senescent cells is their senescence-associated secretory phenotype (SASP) (Basisty *et al*., 2020), a host of secreted factors that are pro-inflammatory and pro-fibrotic. For the FAPs from FID muscles, they displayed many features of senescent cells, including reduction in genes for cell proliferation and enhancement of genes for inflammation (Moiseeva *et al*., 2023), but they lacked upregulation of ECM related genes. The reduction of ECM remodeling and deposition may prevent normal fiber formation during regeneration, as the physical and biochemical cues from the matrix would not be present. Alternatively, this may be an advantage to instances of excess fibrosis, in that the “quasi”-senescent FAPs could not produce increased ECM. Understanding the balance between the beneficial and detrimental consequences of FAPs lacking autocrine IGF-I actions is likely context dependent, and worthy of future investigation.

A larger question that emerges is how the consequences to muscle homeostasis and regeneration differ between the FID model, where FAPs lack IGF-I production, and the complete ablation of FAPs as previously studied (Uezumi *et al*., 2021; Wosczyna *et al*., 2019). The models share regeneration defects, with loss of FAPs having slower regenerative capacity and more negative impact on muscle satellite cell than loss of FAP IGF-I. This suggests that even though the FAPs from FID muscles are defective in many aspects, they must retain some properties that still aid in the differentiation process. While our examination of regenerating muscle spanned the 4 weeks to capture the period in which FAPs are known to contribute to resolving damage, we examined the FAPs most thoroughly at the normal height of expansion, at 3 dpi. Over the course of regeneration, their properties may change and either escape the senescent-like state or adopt more features of senescence. The former possibility did not occur, as senescent FAPs remained following 4 weeks of regeneration in FID muscles. The latter is also unlikely, as one would predict greater ECM accumulation at later timepoints after injury, which we did not observe. The ability for other circulating factors to compensate for the loss of both IGF-I and its signaling pathway members may have helped to resolved muscle damage *in vivo*. For instance, FAPs isolated from FID muscles were still able to respond to TGF-β -induced fibrogenesis and to adipogenic media, albeit more slowly, suggesting that the loss of IGF-I does not block all cytokine mediated communication to the FAPs. Even so, there was still a population of FAPs in FID muscles that bore senescent markers even in non-damaged muscle, as well as a subpopulation of cultured primary FAPs from FID muscles that retained a PDGFRα^+^ state even with fibrogenic or adipogenic signals, which suggests that IGF-I from FAPs is required for them to enter differentiation from quiescence. Taken together, the contribution of other pathways aids in partial rescue of IGF-I dependent deficits that may allow for a phenotype that is milder that that when FAPs are completely absent.

In the current study, we focused on acute muscle regeneration in healthy young adult mice to afford a controlled environment for evaluating the contribution of IGF-I from FAPs on regenerative capacity. During the early stages of muscle regeneration, FAPs interact with satellite cells (SCs) to promote myogenesis (Joe *et al*., 2010). Previous work has shown that FAPs secrete WNT1-inducible signaling pathway protein 1 (WISP1), which regulates SC proliferation by activating the Akt signaling pathway (Lukjanenko *et al*., 2019). Additionally, FAPs produce hepatocyte growth factor (HGF) that may activate quiescent SCs and/or regulate their basal proliferation (Allen *et al*, 1995; Kim *et al*., 2022). Although our RNA-seq data indicated that expression levels of *Wisp1* and *Hgf* were unchanged in the absence of IGF-I, we did not examine the FAP secretome in this study to determine if release of these proteins was affected by the loss of FAP IGF-I. Alternatively, the reduction in SC number could theoretically result solely from a decline in FAP numbers, as FAP ablation also results in reduced SCs in regenerating muscle (Uezumi *et al*., 2021; Wosczyna *et al*., 2019). Although Pax7^+^ SCs are an important part of muscle development and regeneration (Kuang *et al*, 2006; Kuang *et al*., 2007; Relaix *et al*., 2005; von Maltzahn *et al*, 2013), it also requires supporting FAPs for them to thrive in their environment (Joe *et al*., 2010; Kim *et al*., 2022; Lukjanenko *et al*., 2019). If satellite cells acquire deficiencies in their ability to proliferate or differentiate, the consequences will be compounded by the inability of IGF-I deficient FAPs to promote their activation via factors such as HGF (Kim *et al*., 2022). Similarly, a heightened inflammatory environment may go unchecked without the ability for FAPs to interact effectively with the macrophage population. However, the actions of polarized macrophages on FAPs to enhance fatty-fibrotic infiltration in diseased muscles (Liu *et al*, 2023) may also be disrupted by loss of IGF-I from FAPs. Future studies are warranted to determine the potential benefits or disadvantages of removing this source of IGF-I from skeletal muscle. Understanding how the interplay among these cell populations changes when FAPs are in a state of torpor will be important to pursue in light of the reduced ECM deposition in FID muscles, which may be beneficial in disease, but at the expense of reduced regenerative capacity, which may be detrimental. The tissue environment associated with genetic neuromuscular disease or aging may exacerbate the phenotypes we have observed in regenerating FID muscles.

Overall, this study provides multiple lines of evidence that IGF-I produced by FAPs in muscle is a required autocrine factor for FAP actions during muscle regeneration. The consequences of its loss alter FAPs to an extent that they are essentially removed from performing their normal interactions with other cell populations in the muscle, and ultimately stalling regenerative capacity. While IGF-I from FAPs may also have paracrine actions on neighboring cells, these potential direct actions cannot be resolved from those mediated indirectly via its required beneficial actions on FAPs themselves.

## Methods

### Animal Models

All animal experiments were performed under the Institution of Animal Care and Use Committee guidelines previously approved by the University of Florida. Mice were housed under a 12hrs/12hrs day/night cycle and had *ad libitum* access to food and water. To generate the FAP-*Igf1*-Deficient (FID) mice, we crossed Pdgfrα^Cre-ERT2^ mice (Chung et al., 2018) (JAX#032770) with *Igf1^fl/fl^* mice (Liu *et al*, 1998; Spradlin *et al*., 2021) (JAX#012663) and Rosa26^EYFP^ mice (Srinivas *et al*., 2001) (JAX#006148). All animals were maintained on the C57BL/6 background strain. FID mice were homozygous for *Igf1^fl/fl^* and Rosa26^EYFP^, and hemizygous for the *Pdgfra*^Cre-ERT2^ allele since it is homozygous lethal (Kang *et al*, 2010; Kopinke *et al*., 2017; Norris *et al*., 2023). Controls were littermates lacking the *Pdgfra*^Cre-ERT2^ allele. Genotyping was performed in-house as previously described(Spradlin *et al*., 2021). Briefly, 2uL DNA extracted from tails was added into G2 Master mix (M7822, Promega) following manufacturer’s protocol for endpoint PCR, followed by agarose gel electrophoresis. Primers for genotyping are listed in Table 4.

To induce Cre-dependent recombination, Tamoxifen (TAM) dissolved in corn oil was delivered via intraperitoneal (IP) injection to mice at 12-13 weeks of age for 5 consecutive days (100ug/g body weight). five times the amount of body weight (g) in microliters of tamoxifen (TAM) mixed in corn oil (20mg/mL). At least 1 week between the final TAM injection and subsequent experiments was done to allow washout of TAM. Both male and female mice were used as indicated in each experiment.

### Muscle Injury and Tissue Processing

Acute muscle injury was achieved by injection of 50uL PBS containing 1.2% Barium Chloride (BaCl_2_) into the tibialis anterior (TA) muscle in both male and female mice. To induce an adipogenic injury response, 50uL of 50% glycerol in PBS was injected into the TA muscle to induce adipogenic muscle injury. Only females were used for glycerol injections as females generate greater intramuscular adipocytes after glycerol-induced muscle injury (Norris *et al*., 2024; Pisani *et al*, 2010).

Tissues were harvested at 3-28 days-post-injury (dpi) and weighed prior to subsequent steps. To preserve muscles for morphological analysis, muscles were embedded in OCT and then frozen in melting 2-methylbutane cooled by liquid nitrogen. Tissues from glycerol induced muscle injury were performed as previously described (Johnson *et al*., 2022). Muscles used for biochemical analysis were snap-frozen in liquid nitrogen. All samples were stored in −80°C until subsequent measurements.

### Primary Cell Culture

Primary FAPs were isolated from TA muscles at 3dpi. Tissue dissociation followed a protocol that was previously described (Oprescu *et al*, 2020a) with minor adjustments. Briefly, TA muscles were isolated and minced into small pieces followed by 1hr digestion with collagenase II and dispase for 1hr at 37°C on a rotator. Cellular debris were removed using Debris Removal Solution (130-109-398, Miltenyi Biotec) and cells are counted using Countess 3 automated cell counter. Primary FAPs isolation from cell suspensions was achieved using the PDGFRα Microbead Kit coupled with the Magnetic-Activated Cell-Sorting (MACS) system following manufacturer’s protocol (130-101-502, Miltenyi Biotec). Primary FAPs were then cultured onto 24-well plate at 30-35% confluency in proliferation media (PM) (high glucose DMEM, 10% Bovine Calf Serum, 1% Penicillin and Streptomycin, and 1% GlutaMax. The proliferation media was changed every 2-3 days until the cells reached desired confluency. Adipocyte differentiation was performed on cells at 95-100% confluency; Fibroblast differentiation was performed at 65-70% confluency. Adipocyte differentiation was induced with adipocyte differentiation media (DM) as previous described, which contains 5ug/mL insulin (12643-25MG, Sigma) or 10nM of human recombinant IGF-I (100-11-100ug, PeproTech), 5uM of Troglitazone (T2573, Sigma), 3-isobutyl-1-methylxanthine (I5879, Sigma), and Dexamethasone(D4902, Sigma) (Low *et al*, 2017; Norris *et al*., 2023). Fibroblast differentiation was induced with 5ng/mL of TGF-β(AF-100-21C-2ug, PeproTech) as previously described (Contreras *et al*., 2019). For differentiating FAPs, the media was changed daily until the desired time post induction was reached.

### RNA-seq analysis

Total RNA was isolated from primary FAPs isolated from muscles 3 dpi as described above. Reads acquired from the Illumina NovaSeq 6000 platform were cleaned up with the cutadapt program (Martin, 2011) to trim off sequencing adaptors and low-quality bases with a quality phred-like score <20. Reads <40 bases were excluded from RNA-seq analysis. The genome of *Mus musculus* (version GRC38, mm10) from the Ensembl database was used as the reference sequences for RNA-seq analysis. The cleaned reads of each sample were mapped to the reference sequences using the read mapper of the STAR package (Spliced Transcripts Alignment to a Reference, v2.7.9a) (Dobin *et al*, 2013). The mapping results were processed with the HTSeq (High-Throughput Sequence Analysis in Python, v0.11.2)(Anders *et al*, 2015) samtools, and scripts developed in house at ICBR of UF to remove potential PCR duplicates and count uniquely mapped reads for gene expression analysis. PCA analysis (for detecting outlier samples) and volcano plot analysis based on all identified genes in each analysis were performed with the R-package (v4.1.3). The gene expressions were analyzed by a DESeq2-based R pipeline(Love *et al*, 2014). Genes were considered differentially expressed if the log2 fold change was ≥ |1| and if the adjusted *P*-value was ≤ 0.01. Bowtie2 (version 2.5.4) was used for searching gene and exon for each sample(Langmead & Salzberg, 2012).

### Single Cell sample preparation and sequencing

Tissue dissociation was performed following a previously established protocol (Oprescu *et al*., 2020a). For single-cell isolation, ten tibialis anterior (TA) muscles were harvested from five male mice per strain. The muscles were finely minced in wash medium (F-10 supplemented with 10% fetal bovine serum (FBS) and 1% penicillin-streptomycin) to enhance digestion efficiency. Samples were then incubated in dissociation medium, consisting of wash medium supplemented with Collagenase II, Dispase II, and 1% bovine serum albumin (BSA), and rotated at 37°C for 1 hour. Following enzymatic digestion, the suspension was filtered twice through a 40 µm filter to remove residual tendons and debris. Red blood cells were lysed by incubating the suspension in RBC lysis buffer for 3 minutes at room temperature. After obtaining a single-cell suspension, cells were stained with Propidium Iodide (P1304MP, Invitrogen) for 10 minutes to label dead cells, which were subsequently removed using fluorescence-activated cell sorting (FACS) at the Interdisciplinary Center for Biotechnology Research (ICBR) Core, University of Florida. Approximately 200,000 viable cells per sample were collected for downstream analysis. Single-cell RNA sequencing (scRNA-seq) was performed using the 10x Genomics Chromium GEM-X Single Cell 3’ v4 platform, following the manufacturer’s protocol with a targeted recovery of 20,000 cells per sample. Library preparation was conducted according to the commercial protocol, and sequencing was performed on an Illumina NovaSeq X at the ICBR Core. Post-sequencing, reads were aligned and annotated using Cell Ranger (v9.0.0) on HiPerGator (UF Research Computing). On average, ∼40,000 reads per cell were obtained, with approximately 15,000 cells retained per sample after Cell Ranger processing. Downstream analysis was performed primarily using Scanpy (Wolf *et al*, 2018) in Python via Jupyter Notebook.

### Single Cell Analysis: Quality control, dimension reduction

The gene expression matrices from both *Igf1^fl/fl^* and FID samples were merged for joint analysis. Cells with fewer than 200 detected genes and genes expressed in fewer than three cells were excluded. Additionally, cells with mitochondrial gene content exceeding 5% and those expressing more than 6,000 genes were removed. Predicted doublets were identified and excluded using Scrublet (Wolock *et al*, 2019). Each cell was normalized to 10,000 counts per cell to perform unbiased downstream analysis. Samples were then condensed and batch corrected using Harmony (Korsunsky *et al*, 2019). Dimension reduction was performed using Principle Component Analysis (PCA) with 50 neighboring cells.

### DGE, Cell Clustering, Pathway Analysis, and Cell-Cell communication

Cell clusters were identified using differential gene expression (DEG) analysis with the Wilcoxon rank-sum test (p < 0.0001). Each cluster was annotated based on unique cell markers and gene expression profiles using Scanpy. To analyze fibro-adipogenic progenitor (FAP) subclusters, we first extracted FAP-specific clusters, re-normalized gene expression counts to 10,000 counts per cell, and re-clustered the cells. Sub-clustering of FAPs was performed using DEG analysis with Wilcoxon tests (p < 0.0001).

Cell-cell communication and ligand-receptor interactions were analyzed using cell2cell cell2cell(Armingol *et al*, 2022; Dimitrov *et al*, 2022). Gene Ontology (GO) terms was identified using Gene Set Enrichment Analysis (GSEA) and m5 subcollection of GO(m5.go.version2024.1) was utilized for this analysis within FAPs(Smith & Eppig, 2009), and canonical pathway analysis was performed based on m2 (version 2024.1).

### Real-time PCR

Frozen tissue samples were pulverized into a fine powder, and total RNA was isolated using Trizol Reagent (15596026, Thermo Fisher) according to the manufacturer’s protocol. RNA from primary FAPs was extracted with Qiagen RNeasy Micro Kit (74004, Qiagen) following manufacturer’s protocol. RNA samples were then treated with DNase, quantified, and its integrity was verified. Equal quantities of RNA were converted into cDNA using the High-Capacity cDNA Reverse Transcription Kit (4368814, Thermo Fisher). Gene expression levels were assessed in duplicate using the QuantStudio-3 with SYBR Green Master Mix (A46109, Thermo Fisher). The Ct values were normalized against *Gapdh* and/or *Sra-1* as housekeeping genes. *Igf1* deletion PCR was performed as previously described (Spradlin *et al*., 2021). To quantify the extent of deletion using qRT-PCR, amplification of exon3-4 primer pairs of *Igf1*, which includes the deleted sequence of *Igf1*, was normalized to exon6-UTR primer pairs of *Igf1,* which is unaffected by deletion. All primer sequences are listed in Table 4 and schematized in Figure 1C.

### Histological Staining and Analysis

Muscle 8-10 μm cryosections were fixed with 4% Formaldehyde at room temperature (RT) for 10min and then washed in PBS 10min three times. Sections were blocked for 1hr at RT with blocking solution (5% BSA, 0.02% Triton-X100, 0.03%% Sodium Azide). Next, primary antibodies diluted in blocking solution were added onto the samples and then incubated at 4°C overnight. On the second day, samples were washed three times with PBS and then incubated with secondary antibody containing 4’,6-diamidino-2-phenylindole (DAPI) for an 1hr at RT. Samples were then wash three times in PBS containing 0.5% Tween 20 (PBST). Samples were placed at RT until dry, then mounted with Fluoromount-G and cover slips, and sealed with nail-polish. The same procedure was followed for cell culture staining and performed on culture plates or chamber slides. Antibodies are listed in Table 1-2.

**Table 1.** Primary Antibodies.

| Target | Company | Catalog # | Dilution |
| --- | --- | --- | --- |
| Laminin Monoclonal Antibody | Invitrogen | MA1-06100 | 1:1000 |
| Mouse PDGFRa Antibody | R&D system | AF1062 | 1:250 |
| Anti-alpha smooth muscle actin antibody | R&D system | MAB1420 | 1:500~2000 |
| Anti-mouse CD68 antibody | BioLegend | FA-11 | 1:150 |
| Pax-7 Antibody | DSHB | PAX7-S | 1:20 |
| Perilipin-1(D1D8) XP Rabbit mAb | Cell Signaling | 9349 | 1:500 |
| Phospho-Smad2 Rabbit mAb | Cell Signaling | 8828 | 1:1000 |
| Anti-Ki67 Antibody | Abcam | Ab16667 | 1:500 |
| Anti-p53 | Santa Cruz | Sc-126 | 1:100 |
| Anti-IGF1R | Santa Cruz | Sc-271606 | 1:100 |
| Anti-Erk1/2 | Cell Signaling | 4696s | 1:1000 |
| Anti-p-Erk1/2 | Cell Signaling | 9101s | 1:1000 |
| Anti-Akt | Cell Signaling | 2920s | 1:1000 |
| Anti-p-Akt | Cell Signaling | 9271s | 1:1000 |
| Anti-GAPDH | Santa Cruz | Sc-32233 | 1:500 |

**Table 2.** Secondary Antibodies.

| Target | Company | Catalog # | Dilution |
| --- | --- | --- | --- |
| DAPI | Thermo Fisher | D1306 | 1:1000 |
| 555 Donkey Anti-Goat | Thermo Fisher | A32816 | 1:500 |
| 488 Donkey Anti-Rabbit | Thermo Fisher | A-21206 | 1:500 |
| 488 Goat Anti-Rat | Thermo Fisher | A-11006 | 1:500 |
| IRDye 800CW Donkey Anti-Goat | Li-Cor | 926-32214 | 1:15000 |
| IRDye 680 Goat Anti-Mouse | Li-Cor | 926-68070 | 1:15000 |
| IRDye 680 Goat Anti-Rabbit | Li-Cor | 926-68071 | 1:15000 |
| IRDye 800 Goat Anti-Rabbit | Li-Cor | 926-32211 | 1:15000 |

**Table 3.** Reagents.

| Reagent | Company | Catalog # |
| --- | --- | --- |
| G2 Master Mix | Promega | M7882 |
| Tamoxifen | Sigma | T5648-5G |
| Glycerol | Sigma | G5516-500ML |
| Barium Chloride | Sigma | 342920-50G |
| Tissue-Tek O.C.T. Compound | Sakura | 4583 |
| 2-methylbutane | Sigma | 277258-1L |
| Trizol Reagent | Thermo Fisher | 15596026 |
| DNase I | Thermo Fisher | 18047019 |
| High Capacity cDNA Reverse Transcription Kit | Thermo Fisher | 4368814 |
| SYBR Green Master Mix Standard | Thermo Fisher | A46109 |
| RNeasy Micro Kit | Qiagen | 74004 |
| Fluoromount-G | Southern Biotech | 0100-01 |
| Pierce BCA Protein Assay | Thermo Fisher | 23227 |
| NuPAGE Bis-Tris Mini Gel | Thermo Fisher | 10780345 |
| Intercept Blocking Buffer | Licor | 927-60001 |
| Debris Removal Solution | Miltenyi Biotec | 130-109-398 |
| PDGFRa Microbead Kits | Miltenyi Biotec | 130-101-502 |
| IGF-I ELISA Kit | R&D Systems | MG100 |
| Human Insulin | Sigma | 12643-25MG |
| Troglitazone | Sigma | T2573 |
| Dexamethasone | Sigma | D4902-25MG |
| 3-Isobutyl-1-methylxanthine | Sigma | I5879-100MG |
| TGF-β1 | PeproTech | AF-100-21C-2UG |
| Senescence β-Galactosidase Staining Kit | Cell Signaling | 9860S |

**Table 4.**
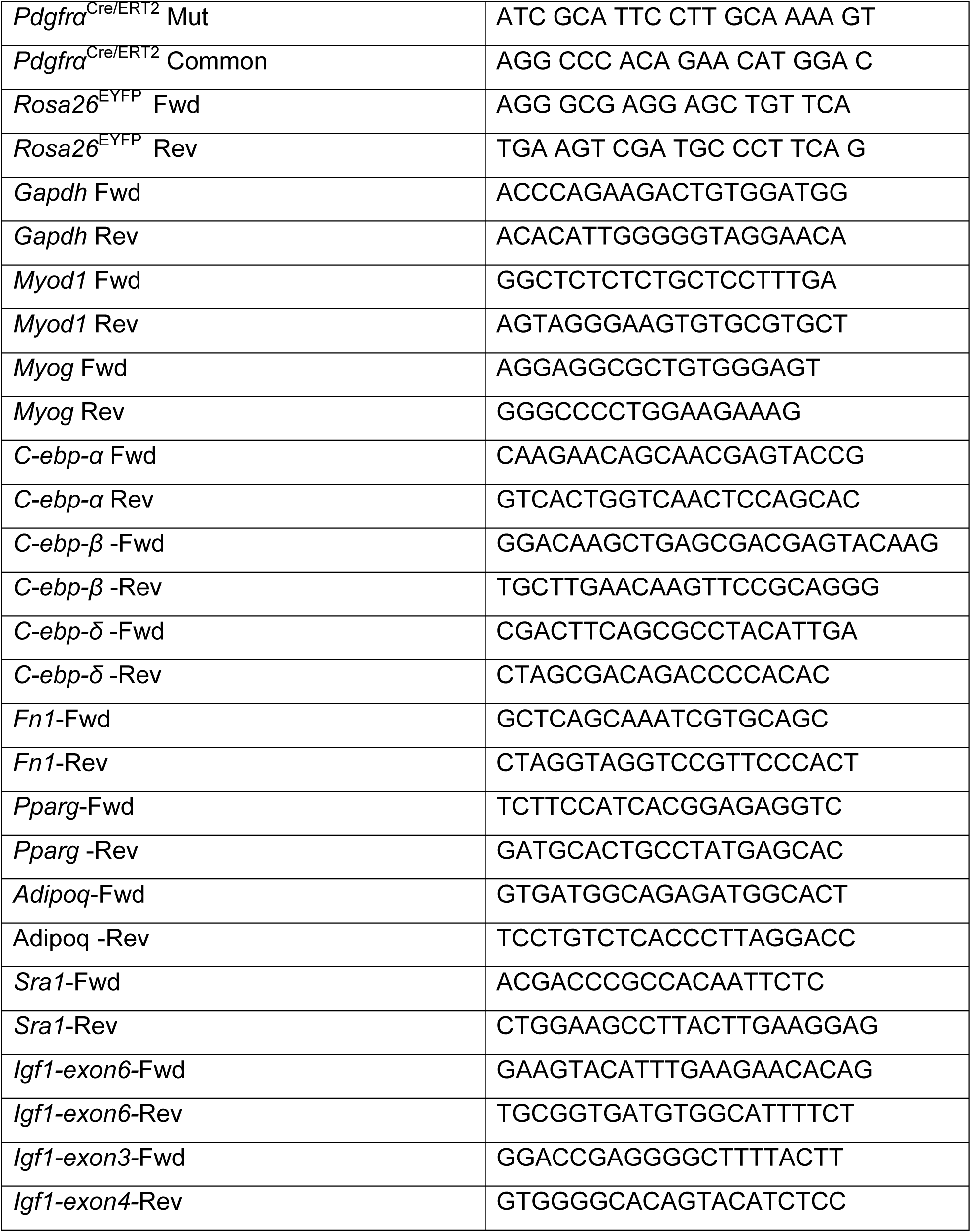
Primer Sequences.

Picro-Sirius Red (PSR) staining was used to determine the total ECM content and fibrosis as previously described without any modification (Smith *et al*, 2020). Muscle cross-sectional areas (CSA) were quantified using SMASH and CellPose (Smith & Barton, 2014; Waisman *et al*, 2021). Quantification of PDGFRα+ and CD68+ cells were performed in ImageJ Software. Images were captured with Keyence X-BZ All-In-One Fluorescent microscope and EVOS Cell Imaging system.

### Immunoblotting

Proteins from tissue and cells were extracted with RIPA buffer (150mM NaCl, 50mM Tris-HCL, 1% Triton-X, 0.5% sodium deoxycholate, 0.1% SDS). The protein concentrations were determined using Pierce BCA Protein Assay (23225, Thermo Fisher) following the manufacturer’s instructions. To prepare sample for western blotting, equal amount of protein were mixed with sample buffer that contains loading dye and 2-mercaptoethanol. After generating the sample mixture, samples were heated at 95°C for 5 mins to denature the proteins. Equal amounts of proteins were loaded onto the 4-12% Bis-Tris Mini gel (NP0321BOX, Invitrogen) and electrophoresis was performed. Proteins were then transferred onto either nitrocellulose or activated PVDF membrane at 60V for 2.5 hr. After transfer, membranes were incubated with Ponceau S (P7170, Sigma) to obtain total protein content and washed by Tris-Buffered Saline (Tween 0.5%) (TBST) to remove Ponceau S. Membranes were incubated in blocking solution (9276001, Licor) for 1hr at RT on shaker. Primary antibodies diluted in blocking solution were added onto the membranes and incubated at 4°C overnight. On the second day, membranes were washed for 10 min three times with TBST. Membranes were then incubated in secondary antibody diluted in blocking solution for 1hr at RT on shaker. Final washes in TBST were performed followed by image capture on Li-Cor Odessey CLX. Quantification of band intensity on membranes was performed in ImageJ studio. Antibodies are listed in Table 1-2.

### IGF-I ELISA

To measure IGF-I levels in the damaged TA muscle at different time point post injury, we used the Quantikine ELISA kit (MG100, R&D Systems) according to the manufacturer’s guidelines. Skeletal muscle lysates were diluted 1:3, and optical density was measured using the SpectraMax M5 Plate Reader (Molecular Devices, Sunnyvale, CA) at 450 and 570 nm. Wavelength correction was performed by subtracting the 570 nm readings from those at 450 nm. IGF-I concentrations were calculated from the optical density of the averaged duplicates using the 4-Parameter standard curve provided by the plate reader. Muscle IGF-I levels were normalized to protein concentration, measured with the Pierce BCA Protein Assay Kit (23227, Thermo Fisher). Results are expressed as picograms IGF-I per microgram total protein.

### Statistics

Comparisons were performed by unpaired Students t-test, one-way and two-way ANOVAs, following by post hoc testing where appropriate. P-values < 0.05 were considered significant.

## Acknowledgments

This work was supported by the Spinal Muscular Atrophy Foundation to ERB, and by US National Institute of Health (NIH) P50 AR052646 to ERB supporting the Physiological Assessment Core. The authors acknowledge University of Florida Research Computing for providing computational resources and support that have contributed to the research results reported in this publication

## Author Contributions

YEL, DK, FY, and ERB conceived and designed the study. YEL, ZA-T, TYA, L-YK, FY, YZ, BES, and ABA performed all experiments. All authors analyzed the data. YEL and ERB wrote the manuscript.

## Data Availability Statement

The data that support the findings of this study are available in the methods and/or results of this article. All software and instrumentation have been described within the manuscript text.

## Supplemental Information

Figures S1-S10.

